# Heat hardening enhances mosquito heat tolerance in a species-specific and trait-specific manner

**DOI:** 10.1101/2025.09.14.676167

**Authors:** Apeksha L. Warusawithana, Belinda van Heerwaarden, Ary A. Hoffmann, Perran A. Ross

## Abstract

Models predict that the distribution of ectotherms including mosquitoes will shift with climate change, but few incorporate adaptive capacity. Acclimation is one mechanism by which mosquitoes could adapt, allowing mosquitoes that have experienced sub-lethal stress previously to tolerate subsequent stressful environments. In this study we evaluated the heat tolerance of three vector mosquito species, *Aedes aegypti*, *Ae. notoscriptus* and *Culex quinquefasciatus,* after being previously exposed to heat hardening. Adult males and females were heat-hardened by exposure to 41°C for one hour and subsequently tested for heat survival and knockdown following one-hour heat shocks across a range of temperatures up to the lethal limit, ramping CTmax assays and static temperature knockdown time assays. The three species differed markedly in their heat tolerance across all assays, with *Ae. aegypti* being the most heat tolerant and *Cx. quinquefasciatus* being the least. Females from all three species were more heat tolerant than males in the one-hour heat shock assays, but effects of sex were absent or inconsistent for CTmax and heat knockdown time assays. A beneficial impact of heat hardening on subsequent heat shock knockdown was evident in both sexes of all three species. However, hardening effects differed substantially for survival 24 hr later, ranging from no effect of hardening in *Cx. quinquefasciatus* to a ∼1°C increase in LT_50_ in *Ae. notoscriptus*. In contrast, no effects of heat hardening were detected for CTmax or static knockdown time assays. An additional experiment in *Ae. aegypti* detected no benefits of heat shock exposure in female patents on the thermal tolerance of offspring. Our findings emphasize the need to consider effects of acclimation including heat hardening in models to predict the response of mosquitoes to climate warming. They also have implications for measuring thermal tolerance in mosquitoes more generally, given that both sex and hardening effects depend on the type of assay used and trait measured.

## Introduction

Climatic change is a key factor reshaping ecosystems, thereby altering species’ distributions and dynamics across the globe (Grimm et al., 2013; Scheffers et al., 2016). Mean global temperatures have already surpassed the historical target of 1.5 °C of warming (Cointe & Guillemot, 2023), reaching 1.55 °C in 2024 (WMO, 2025). Ectothermic insects, including mosquitoes, are expected to be significantly affected by environmental warming, as their physiological and biological traits are directly influenced by ambient temperature and by temperature extremes (Carrington. et al., 2013; Harvey et al., 2020; Hoffmann et al., 2003), ultimately determining their overall fitness (Ciota et al., 2014; Cossins & Bowler, 2012; Lahondere & Bonizzoni, 2022).

While insects are vulnerable to high temperatures (Ma et al., 2021), mechanisms related to plasticity may allow them to cope with extreme thermal conditions (Rodrigues & Beldade, 2020; Sgro et al., 2016), which we consider here as the ability of a genotype to express different phenotypes in response to environmental stimuli (Lalejini et al., 2021; Rodrigues & Beldade, 2020; West-Eberhard, 1989). Plasticity allows organisms to respond to short term environmental changes more rapidly than through evolutionary adaptation alone (Couper et al., 2021; Kellermann & van Heerwaarden, 2019).

Mosquitoes inhabit highly dynamic environments including temperate regions characterised by seasonality, where phenotypic plasticity may expand the conditions under which mosquitoes can maintain their performance (Bujan et al., 2020; Urbanski et al., 2010). For example, populations of the Asian tiger mosquito, *Aedes albopictus,* from temperate regions exhibit seasonal phenotypic variation in egg surface hydrocarbon composition, whereas tropical populations do not (Urbanski et al., 2010). Similarly, Oliveira et al. (2021) found that CTmin in *Aedes* and *Culex* mosquitoes tracked seasonal variation in a temperate climate, but not in a tropical climate. However, plasticity does not always enhance thermal performance; depending on the intensity and duration of exposure, it can also cause physiological costs (Arnold et al., 2019; Gunderson et al., 2017; Hoffmann & Bridle, 2022; Watkins, 2021).

Heat hardening, a short-term form of phenotypic plasticity, occurs when prior exposure to sublethal high temperatures enhances subsequent thermal performance (Bowler, 2005; Hoffmann et al., 2023; Sørensen et al., 2016). While this response has been widely studied in some insects and particularly in *Drosophila melanogaster* (Ferguson et al., 2024; Kellermann & Sgrò, 2018; Sejerkilde et al., 2003; Sørensen et al., 2016), it remains understudied in mosquitoes. In the context of predicting the impact of global warming on mosquitoes, it is important to investigate whether mosquitoes employ heat hardening as a rapid response mechanism to enhance their thermal performance under stressful temperatures. Additionally, it is important to understand whether parental heat stress exposure can induce transgenerational plasticity, allowing offspring to rapidly respond to environmental changes without underlying genetic alterations (Foo et al., 2019; Fox & Mousseau, 1998).

Consequently, understanding the capacity of mosquitoes to undergo heat hardening is crucial for predicting both their ability to sustain populations and their potential to transmit pathogens under climate change. To maintain mosquito populations, adult females must survive long enough to complete at least one gonotrophic cycle, from blood feeding to oviposition (Brady et al., 2013). Importantly, in the context of pathogen transmission, infected female mosquitoes must survive long enough for the pathogen to complete its extrinsic incubation period and become infectious (de Souza & Weaver, 2024; Kamiya et al., 2020; Mordecai et al., 2019; Reeves et al., 1994), and for subsequent blood feeding to occur, enabling transmission to an uninfected host (Barr et al., 2024; Goindin et al., 2015; Muturi et al., 2008). Hence, mosquito survival is a critical determinant of vectorial capacity, raising the question of whether heat hardening can enhance survival under subsequent heat stress to allow completion of one or more gonotrophic cycles.

Although numerous studies have examined how vector mosquitoes respond to temperature (Carlassara et al., 2024; Chura et al., 2023; Colinet et al., 2015; Delatte et al., 2009), relatively few have addressed their capacity for plasticity to buffer stressful high temperatures (Ciota et al., 2014; Gray, 2013; Sivan et al., 2020). Most studies have focused on developmental plasticity, assessing adult trait performance after rearing immature stages at elevated constant temperatures, which can reduce adult mosquito survival (Ciota et al., 2014), body size (Ciota et al., 2014), fecundity (Ezeakacha & Yee, 2019; Pekľanská et al., 2025), and egg hatch (Pekľanská et al., 2025). Despite well documented negative effects, the ability of thermal hardening to alleviate them has rarely been considered. Sivan et al. (2020) found that larval pre-exposure to sublethal heat shock improved *Ae. aegypti* larval survival following subsequent heat stress.

This study was designed to comprehensively examine the effect of adult heat hardening on heat tolerance in mosquitoes, including transgenerational impacts. We investigate the effect of heat hardening (through a sublethal one-hour heat shock at 41 °C) on heat tolerance in three vector mosquito species, *Aedes aegypti, Aedes. notoscriptus,* and *Culex quinquefasciatus. Aedes aegypti* is a major vector of dengue (Brady & Hay, 2020; Jansen & Beebe, 2010), placing over 3.9 billion people in more than 132 countries at risk, with approximately 96 million symptomatic cases and 40, 000 deaths reported annually (WHO, 2025). *Aedes notoscriptus* is a key vector of Ross River fever (Claflin & Webb, 2015), responsible for around 5000 human infections annually in Australia (AGDH, 2025), and recently implicated as a vector of *Mycobacterium ulcerans* in south-eastern Australia (Mee et al., 2024). According to the World Health Organization (WHO, 2025), lymphatic filariasis, primarily transmitted by *Cx. quinquefasciatus* (Negi & Verma, 2018), remains a public health threat despite the launch of the WHO’s Global Programme to Eliminate Lymphatic Filariasis (GPELF) in 2000, with over 657 million people in 39 countries still at risk worldwide. By considering these three species we aim to develop generalizations around hardening across mosquitoes. We also consider transgenerational effects in one of these species.

## Methodology

### Mosquito colonies and rearing

*Aedes aegypti*, *Ae. notoscriptus* and *Cx. quinquefasciatus* colonies were originally established from eggs collected in Queensland, Australia, and have been maintained in the laboratory for at least 50 generations. Mosquitoes were reared under standard mosquito insectary conditions of 26 ± 0.5 °C and 50-70% relative humidity (RH), with a 12:12 h (light: dark) photoperiod, following the protocol described in Ross et al. (2017) with modifications. Adults were fed with a 10% sucrose solution that was replaced with water one day before blood feeding. Females were blood fed on the forearm of a single adult human volunteer which was approved by the University of Melbourne Human Ethics committee (project ID 28583). *Aedes aegypti* and *Ae. notoscriptus* eggs were collected by lining a plastic cup (500 mL) containing 250 mL of larval rearing water with a strip of sandpaper (10 cm x 4 cm, Norton Master Painters P80; Saint-Gobain Abrasives Pty. Ltd., Thomastown, Victoria, Australia) while *Cx. quinquefasciatus* egg rafts were collected from the surface of the water.

*Aedes aegypti* and *Ae. notoscriptus* eggs (< 1 week old) were induced to hatch by adding 5-10 grains of active dry yeast in reverse osmosis (RO) water (500 mL), while *Cx. quinquefasciatus* eggs were left to hatch and larvae were reared in hay-infused (for at least one week) RO water. The larval density for all three species was controlled to approximately 500 larvae per plastic tray (40 cm x 30 cm x 6.5 cm) with 4 L of water. We provided 0.05 mg of food (Hikari tropical sinking wafers, Kyorin food, Himeji, Japan) per larva per day. Pupae were separated by sex based on their body size and shape and the sex of adults was confirmed within 24 hours of emerging. Adult mosquito density was maintained at approximately 500 per cage (27 L BugDorm-1, Megaview Science C Ltd., Taichung, Taiwan).

### Heat hardening treatment

Heat hardening of adult *Ae. aegypti, Ae. notoscriptus,* and *Cx. quinquefasciatus* involved placing a maximum of 100 mosquitoes per small plastic cage (1.5 L) in a pre-programmed incubator (PHcbi MIR-254) at 41 °C in the presence of 2-3 water-soaked cotton balls (to prevent desiccation stress) for one hour. Mosquitoes were then returned to 26° C and allowed to recover for 24 hr. This hardening treatment was based on a pilot experiment, with 41 °C being the maximum temperature with a 100% survival and 0% knockdown across the three species. Basal mosquitoes were derived from the same cohort as hardened mosquitoes but were not exposed to the heat hardening treatment (Figure 1).

**Figure 1.**
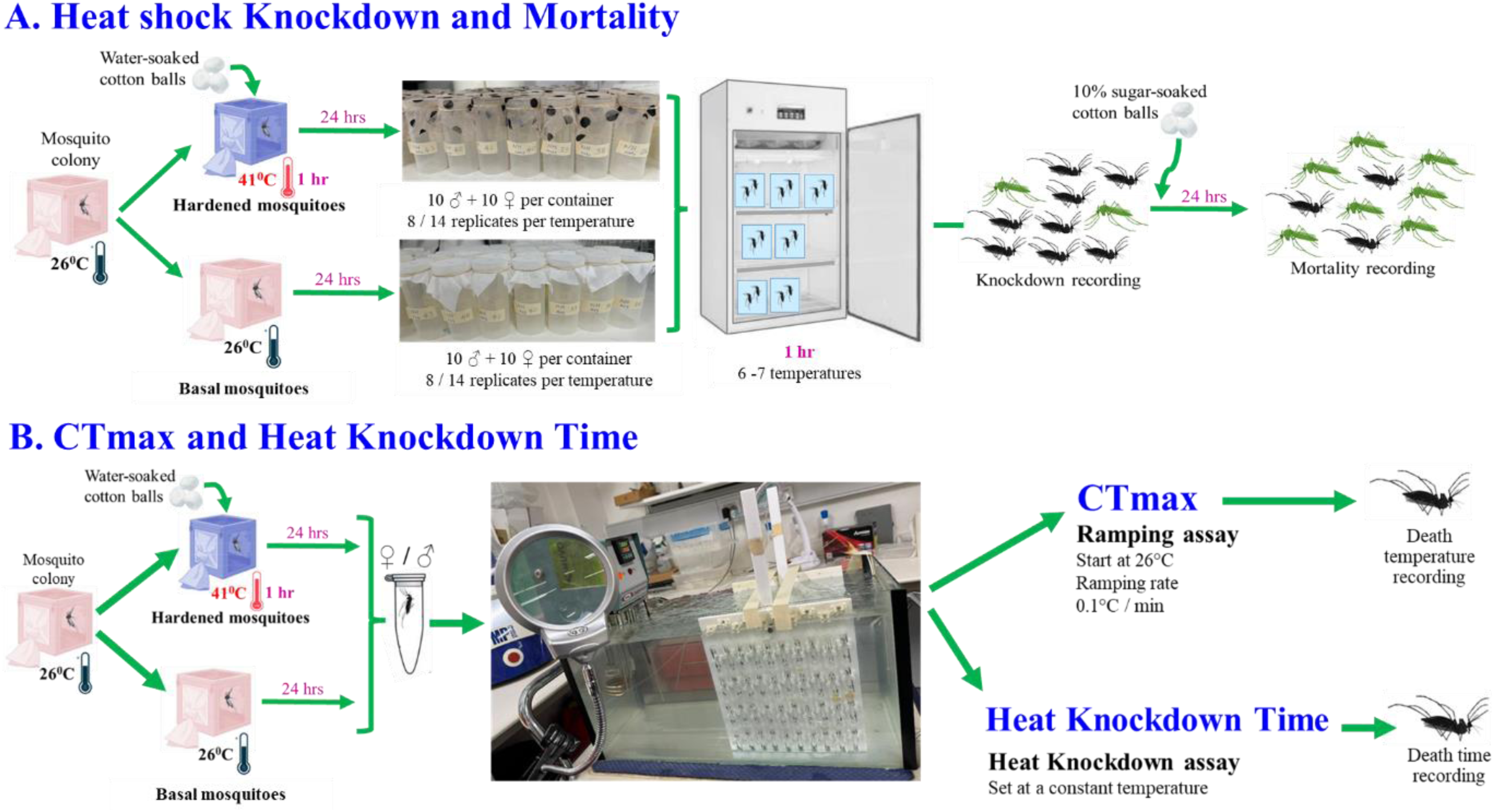
Experimental design: A. Heat shock knockdown and mortality: Individuals were exposed to a heat hardening treatment (one-hour heat shock at 41 °C), followed by a 24-hour recovery period. They were then exposed to a range of constant temperatures (6-7temperatures) for one hour. Heat knockdown was recorded immediately after exposure and mortality was recorded 24 hours later; B. CTmax and heat knockdown time: Individuals were exposed to the same heat hardening treatment (one-hour heat shock at 41°C), followed by a 24-hour recovery period. CTmax was measured using a ramping assay, and heat knockdown time was recorded using a heat knockdown assay. The same procedures were applied to offspring in cross-generational experiments, where the parental generation received the same one-hour heat shock at 41°C.

### Heat hardening effects on heat shock knockdown and mortality

To determine the effects of heat hardening on heat shock knockdown and mortality, hardened and basal male and female adults (4-5 days old) of each species were exposed to heat shocks at 6 – 7 constant temperatures in pre-programmed incubators (PHcbi MIR-254). Adults were placed in plastic cups (125 mL) covered with a fine mesh with no access to water and exposed to one-hour heat shocks in 1 °C increments between 38 – 44 °C, 38 – 43 °C, and 36 – 42 °C for *Ae. aegypti, Ae. notoscriptus,* and *Cx. quinquefasciatus* respectively. These temperatures were chosen based on pilot experiments. Water-soaked cotton was not provided as this could have inadvertently affected mortality by wetting and impairing the mosquitoes. For each temperature treatment, a total of 14 replicates were conducted for *Ae. aegypti* and *Ae. notoscriptus,* and 8 replicates for *Cx. quinquefasciatus*, with each replicate consisting of 10 males and 10 females in a cup. Concurrently, for each species and hardening treatment, an untreated control group was maintained at 26 °C, consisting of 3 replicates, each with 10 males and 10 females. Knockdown was scored immediately after exposure to each temperature by counting the number of males and females in each cup that were knocked down and unable to right themselves. Mosquitoes were then provided with water and sugar, and mortality was recorded 24 hours after exposure to each temperature (Figure 1A).

### Heat hardening effects on critical thermal maxima (CTmax)

A thermal ramping assay was followed to measure the effects of heat hardening on CTmax (Jørgensen et al., 2019; MacLean et al., 2019; Overgaard et al., 2012). Hardened and basal groups of male and female mosquitoes (5-9 days old) from all three species were transferred individually into pre-numbered 5 mL glass vials with plastic screw tops. Two racks, each holding 40 randomly mounted vials, were submerged in a glass tank (60 cm x 30 cm x 30 cm) filled with water set to 26 °C using a thermoregulator (Ratek Instruments, Boronia, Victoria, Australia). Before the racks were submerged, the water tank was programmed to gradually increase the water temperature at a rate of 0.1 °C per minute. Each mosquito was closely monitored to record the water temperature at which no movement was detectable. The water in the tank was continuously stirred by a pump to ensure water temperature homogeneity and monitored with a digital temperature probe. Sixty mosquitoes per treatment group (hardened and basal), per sex (♀ and ♂), and per species (*Ae. aegypti, Ae. notoscriptus,* and *Cx. quinquefasciatus*) were tested. The experiment was conducted in three blocks across three age groups (5, 6 and 9 days old) for *Ae. aegypti* and *Ae. notoscriptus*, and in four blocks across four age groups for *Cx. quinquefasciatus* (5, 7, 8 and 9 days old) (Figure 1B).

### Heat hardening effects on static heat knockdown time

To determine the effects of heat hardening on static heat knockdown time, mosquitoes were exposed to a thermal knockdown time assay (Dennington et al., 2024; Jorgensen et al., 2021; Jørgensen et al., 2019) at constant temperatures of 43.1 °C, 41.2 °C, or 39.8 °C for *Aedes aegypti, Ae. notoscriptus,* and *Cx. quinquefasciatus* respectively. Temperatures had been selected based on the mean CTmax values obtained in the previous experiment, i.e., 43.6 °C, 41.5 °C, and 40.0 °C, respectively. The temperatures used did not precisely match CTmax, reflecting minor differences between the set and actual water bath temperatures. The mosquitoes were exposed and scored according to the previous experiment by recording the time at which a mosquito ceased movement. Sixty mosquitoes per treatment group, per sex, and per species were tested in 3-4 blocks with 3-4 age groups, consistent with the previous experiment (Figure 1B).

### Cross-generational effects of heat shock exposure in *Aedes aegypti*

To test for cross-generational effects of heat shock exposure in *Ae. aegypti*, mated parental female mosquitoes (4-5 days old) were exposed to a one-hour heat shock at 41 °C without access to water (i.e., parental exposure). Afterward, the mosquitoes were provided with a 10% sugar solution and maintained under standard insectary conditions for up to 24 hours to allow recovery. Both treated and untreated females were blood-fed, their eggs were collected, and their offspring were reared under standard conditions to obtain adults for subsequent experiments. Male and female offspring (4-5 days old) from treated or untreated female parents were exposed to a heat shock survival assay at temperatures ranging from 38 °C to 45 °C, following the experimental procedure described above, with each treatment having 8 replicates (Figure 1A). For the CTmax and heat knockdown time experiments conducted on offspring (5-9 days old) of treated or untreated female parents, the same procedures and replication levels were applied as described above (Figure 1B).

### Data analysis

All statistical analysis and data visualisation were performed using R in RStudio (version 2024.04.01+748; (RStudio Team, 2024). The core packages used included drc (version 3.0.1; Ritz et al., 2015), tidyverse (version 2.0.0; Wickham et al., 2019), ggplot2 (version 3.5.2; Wickham, 2016), ggpattern (version 1.0.1; Mike et al., 2023), car (version 3.1.2; Fox and Weisberg, 2023), dplyr (version 1.1.4; Wickham et al., 2023), lme4 (version 1.1.35.3; Bates et al., 2015), gridExtra (version 2.3; Auguie and Antonov, 2017), and Hmisc (version 5.1.2; Harrell and Dupont, 2023). Statistical significance was evaluated using the 0.05 significance level and variation in parameter estimates are shown through standard errors and 95% confidence intervals.

#### Dose-response modelling for knockdown and mortality

Heat shock knockdown and mortality were analysed separately for three mosquito species: *Ae. aegypti*, *Ae. notoscriptus*, and *Cx. quinquefasciatus.* For each species, dose-response curves were fitted using the drm() function from the drc package (version 3.0.1; Ritz et al. 2015). The number of affected individuals (knocked down and dead) at each experimental temperature was modelled using three-parameter log-logistic model (LL.3). The treatment factor included combinations of thermal exposure conditions (basal or hardened) and sex (♀ or ♂) in 6-7 temperature groups, assuming a binomial distribution.

#### Estimation and comparison of median lethal temperature (LT_50_)

The median lethal temperature (LT_50_) defined as the temperature at which 50% of individuals were knocked down or died, was estimated for each treatment group using the effective dose function, ED (), with delta method-based confidence intervals, which allowed quantification of thermal tolerance for both knockdown and mortality endpoints across the different species and treatment groups. ED refers to the estimated LT_50_ values from the dose-response model. To assess statistical differences in LT_50_ values within each species, the EDcomp () function from the drc package (version 3.0.1; (Ritz et al., 2015) was used. This function performs pairwise comparisons of effective dose (LT_50_) between groups and provides ANOVA-style significance testing and the resulting *p*-values were used to identify meaningful differences between treatments or sexes.

#### Critical thermal maximum (CTmax) and heat knockdown time

A general linear model was used to assess the main and interactive effects of sex, age, and hardening treatment on CTmax and heat knockdown time, treating each species separately. The age variable was treated as a categorical variable for group differences. A full interaction model structure was used to include relevant two-way and three-way interactions. Model assumptions were checked using residual and Q-Q plots. Statistical significance of main and interaction terms was assessed through Type III ANOVAs via the Anova () function from the car package (version 3.1.2; (Fox & Weisberg, 2023).

#### Figure visualization

Dose-response curves for both knockdown and mortality were visualised using base R plotting functions. All the boxplots for CTmax and heat knockdown time were generated using ggplot2 (version 3.5.2; (Wickham, 2016) and enhanced with ggpattern (version 1.0.1; (Mike et al., 2023) for visual differentiation of treatment groups. To enable visual comparisons across species and traits, multi-panel composite figures were generated using gridExtra package (version 2.3; (Auguie & Antonov, 2017).

## Results

### Heat hardening effects on heat shock knockdown and mortality

The effect of heat hardening (one-hour heat shock at 41 °C) on heat knockdown (Figure 2A-C, Table 1) and mortality (Figure 2D-F, Table 2) was assessed in male and female mosquitoes from three species (*Ae. aegypti*, *Ae. notoscriptus*, and *Cx. quinquefasciatus*). *Ae. aegypti* was the most heat-resistant species, followed by *Ae. notoscriptus,* while *Cx. quinquefasciatus* was the least tolerant both in terms of knockdown and mortality (Figure 2). Across all species, sexes and treatments, mortality LT_50_ values were approximately 2 °C higher than knockdown LT_50_ values (Tables 1-2), suggesting that many individuals knocked down recovered within 24 hours following heat exposure. No knockdown or mortality was observed in any of the basal or hardened controls.

**Figure 2.**
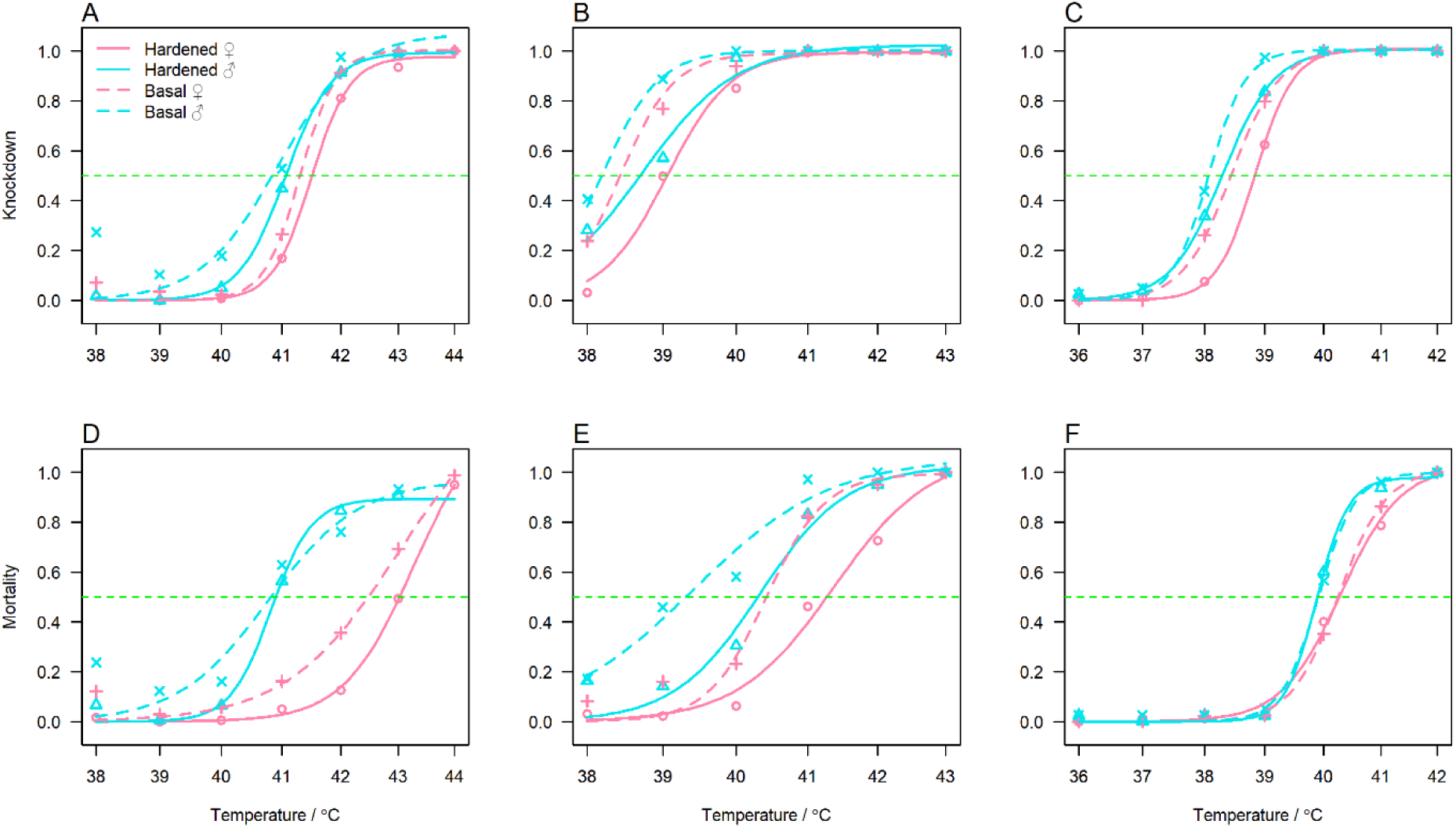
Heat knockdown (A, B, C) and 24-hr mortality (D, E, F) curves for basal and hardened *Ae. aegypti* (A, D), *Ae. notoscriptus* (B, E), and *Cx. quinquefasciatus* (C, F) males and females. Symbols represent the mean percent knockdown or mortality averaged across 8 (*Cx. quinquefasciatus*) or 14 (*Ae. aegypti* and *Ae. notoscriptus*) replicates per treatment, sex and temperature. Lines represent the predicted probability of knockdown or mortality across exposed temperatures: pink lines represent females, blue lines indicate males, solid lines indicate hardened mosquitoes, and dashed lines indicate basal mosquitoes. Intersections with green lines represent LT_50_ for knockdown and mortality.

**Table 1.** Estimated LT_50_ (°C) for heat knockdown of *Ae. aegypti*, *Ae. notoscriptus*, and *Cx. quinquefasciatus* following a one-hour heat shock.

|  | LT<sub>50</sub> | Std. Error | Lower 95% CI | Upper 95% CI |
| --- | --- | --- | --- | --- |
| *Ae. aegypti* |  |  |  |  |
| Basal ♀ | 41.305 | 0.057 | 41.193 | 41.416 |
| Basal ♂ | 40.924 | 0.088 | 40.750 | 41.097 |
| Hardened ♀ | 41.494 | 0.058 | 41.380 | 41.608 |
| Hardened ♂ | 41.061 | 0.060 | 40.943 | 41.179 |
| *Ae. notoscriptus* |  |  |  |  |
| Basal ♀ | 38.426 | 0.084 | 38.260 | 38.592 |
| Basal ♂ | 38.189 | 0.082 | 38.028 | 38.349 |
| Hardened ♀ | 39.055 | 0.082 | 38.895 | 39.216 |
| Hardened ♂ | 38.726 | 0.096 | 38.538 | 38.914 |
| *Cx. quinquefasciatus* |  |  |  |  |
| Basal ♀ | 38.440 | 0.049 | 38.343 | 38.537 |
| Basal ♂ | 38.072 | 0.036 | 38.002 | 38.142 |
| Hardened ♀ | 38.837 | 0.040 | 38.758 | 38.916 |
| Hardened ♂ | 38.296 | 0.049 | 38.198 | 38.393 |

**Table 2.**
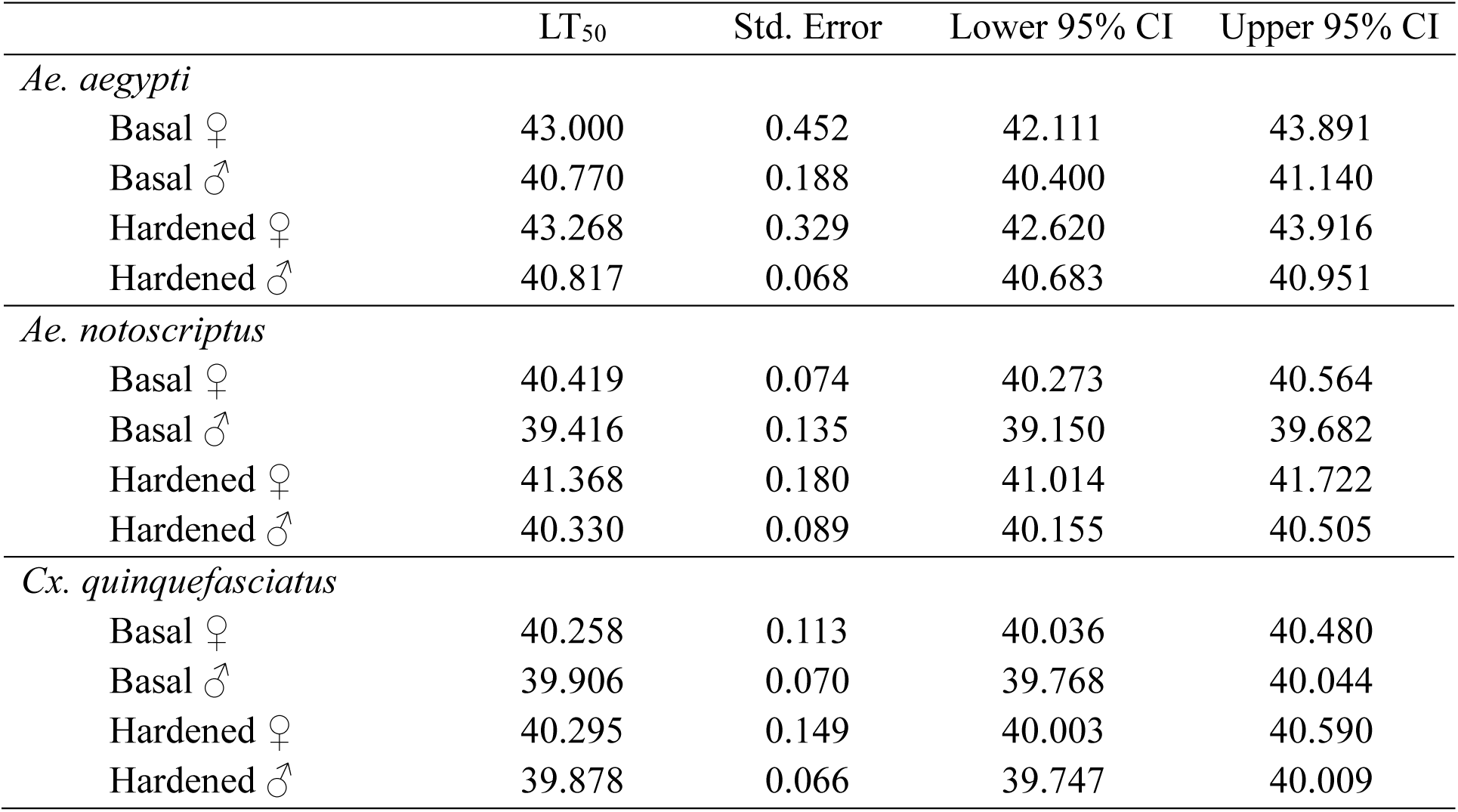
Estimated LT_50_ (°C) for 24-hr mortality of *Ae. aegypti*, *Ae. notoscriptus*, and *Cx. quinquefasciatus* following a one-hour heat shock.

|  | LT<sub>50</sub> | Std. Error | Lower 95% CI | Upper 95% CI |
| --- | --- | --- | --- | --- |
| *Ae. aegypti* |  |  |  |  |
| Basal ♀ | 43.000 | 0.452 | 42.111 | 43.891 |
| Basal ♂ | 40.770 | 0.188 | 40.400 | 41.140 |
| Hardened ♀ | 43.268 | 0.329 | 42.620 | 43.916 |
| Hardened ♂ | 40.817 | 0.068 | 40.683 | 40.951 |
| *Ae. notoscriptus* |  |  |  |  |
| Basal ♀ | 40.419 | 0.074 | 40.273 | 40.564 |
| Basal ♂ | 39.416 | 0.135 | 39.150 | 39.682 |
| Hardened ♀ | 41.368 | 0.180 | 41.014 | 41.722 |
| Hardened ♂ | 40.330 | 0.089 | 40.155 | 40.505 |
| *Cx. quinquefasciatus* |  |  |  |  |
| Basal ♀ | 40.258 | 0.113 | 40.036 | 40.480 |
| Basal ♂ | 39.906 | 0.070 | 39.768 | 40.044 |
| Hardened ♀ | 40.295 | 0.149 | 40.003 | 40.590 |
| Hardened ♂ | 39.878 | 0.066 | 39.747 | 40.009 |

In all three species, the knockdown and mortality curves indicated greater heat tolerance in females compared to males, regardless of hardening treatment (Figure 2). Basal females exhibited significantly higher knockdown tolerance than basal males in all three species, with the greatest difference observed in *Ae. aegypti* (*Ae. aegypti* ΔLT_50_ = 0.381 °C, *t*_18_ = 3.615, *p* < 0.001; *Cx. quinquefasciatus* ΔLT_50_ = 0.368 °C, *t*_18_ = 6.039, *p* < 0.001; *Ae. notoscriptus* ΔLT_50_ = 0.238 °C, *t*_18_ = 2.015, *p* = 0.048; Tables 1, S1). Basal females also had significantly higher heat tolerance than basal males in terms of mortality, with sex differences in *Ae. aegypti* being especially pronounced (*Ae. aegypti* ΔLT_50_ = 2.231 °C; *Ae. notoscriptus* ΔLT_50_ = 1.003 °C; *Cx. quinquefasciatus* ΔLT_50_ = 0.352 °C, all *p* < 0.009; Tables 2, S2). The higher heat tolerance of females was maintained even after the hardening treatment in all three species, both in terms of knockdown (*Ae. aegypti* ΔLT_50_ = 0.433 °C; *Ae. notoscriptus* ΔLT_50_ = 0.329 °C; *Cx. quinquefasciatus* ΔLT_50_ = 0.540 °C, all *p* < 0.009; Tables 1, S1) and mortality (*Ae. aegypti* ΔLT_50_ = 2.451 °C; *Ae. notoscriptus* ΔLT_50_ = 1.038 °C; *Cx. quinquefasciatus* ΔLT_50_ = 0.418 °C, all *p* < 0.011; Tables 2, S2).

Overall, heat hardening increased the estimated LT_50_ values compared to the basal counterparts, though effects varied across the species, sexes and traits (Figure 2). Hardening effects were strongest in *Ae. notoscriptus*, with heat hardening increasing the knockdown LT_50_ by 0.629 °C (*t*_18_ = - 5.405, *p* < 0.001) in females and 0.538 °C (*t*_18_ = - 4.311, *p* < 0.001) in males (Tables 1, S1). Effects were even more substantial for mortality, with hardening increasing the mortality LT_50_ by 0.949 °C (*t*_18_ = - 4.983, *p* < 0.001) in females and 0.914 °C (*t*_18_ = - 5.685, *p* < 0.001) in males (Tables 2, S2). Both *Ae. aegypti* and *Cx. quinquefasciatus* benefitted less from heat hardening compared to *Ae. notoscriptus* across both traits. Although hardening improved knockdown LT_50_ values for *Cx. quinquefasciatus* (females: ΔLT_50_ = 0.396 °C, *t*_18_ = - 6.271, *p* < 0.001; males: ΔLT_50_ = 0.224 °C, *t*_18_ = - 3.687, *p* < 0.001; Tables 1, S1), mortality was not significantly affected by heat hardening in either sex (females: *t*_18_ = - 0.204, *p* = 0.839; males: *t*_18_ = 0.286, *p* = 0.775; Tables 2, S2). In *Ae. aegypti,* although LT_50_ values for both knockdown (Δ LT_50_ ♀ = 0.189 °C; Δ LT_50_ ♂ = 0.137 °C; Tables 1, S1) and mortality (Δ LT_50_ ♀ = 0.267 °C; Δ LT_50_ ♂ = 0.047 °C; Tables 2, S2) tended to increase after heat hardening, none of these differences were statistically significant (all *p* > 0.198) except for knockdown tolerance in females (*t*_18_ = - 2.340, *p* = 0.020; Tables 1, S1).

### Heat hardening effects on critical thermal maxima (CTmax)

The effect of heat hardening (one-hour heat shock at 41 °C) on critical thermal maxima (CTmax) was assessed in male and female mosquitoes from the three species (Figure 3). Clear inter-species differences in CTmax were evident, with *Ae. aegypti* exhibiting a higher estimated CTmax (44.378 °C) than *Ae. notoscriptus* (42.098 °C) and *Cx. quinquefasciatus* (40.767 °C). Heat hardening increased CTmax by 0.023 °C in *Ae. notoscriptus* and by 0.055 °C in *Cx. quinquefasciatus* but decreased it by 0.037 °C in *Ae. aegypti.* However, none of these effects were statistically significant (all *p*-values > 0.302; Table S3). A sex difference in heat tolerance was detected in *Cx. quinquefasciatus,* with males showing lower CTmax than females (Δ = - 0.121, *F*_1, 234_ = 5.219, *p* = 0.023). In contrast, males exhibited higher tolerance than females in both *Ae. aegypti* (Δ = 0.258, *F*_1, 234_ = 8.912, *p* = 0.003) and *Ae. notoscriptus* (Δ = 0.057, *F*_1, 234_ = 1.268, *p* = 0.261), but differences were only statistically significant in *Ae. aegypti* (Table S3). CTmax tended to decrease in later blocks where mosquitoes were older, with a significant negative effect of age detected across species (all *p*-values < 0.001; Table S3). On average, CTmax decreased by approximately 0.1 °C for each day of age across species (Table S3). When considering interaction effects among sex, hardening treatment and age across all three species, none of the two-way or three-way interactions significantly affected CTmax (all *p*-values > 0.424; Table S3), indicating that none of the examined factors interacted to influence CTmax.

**Figure 3.**
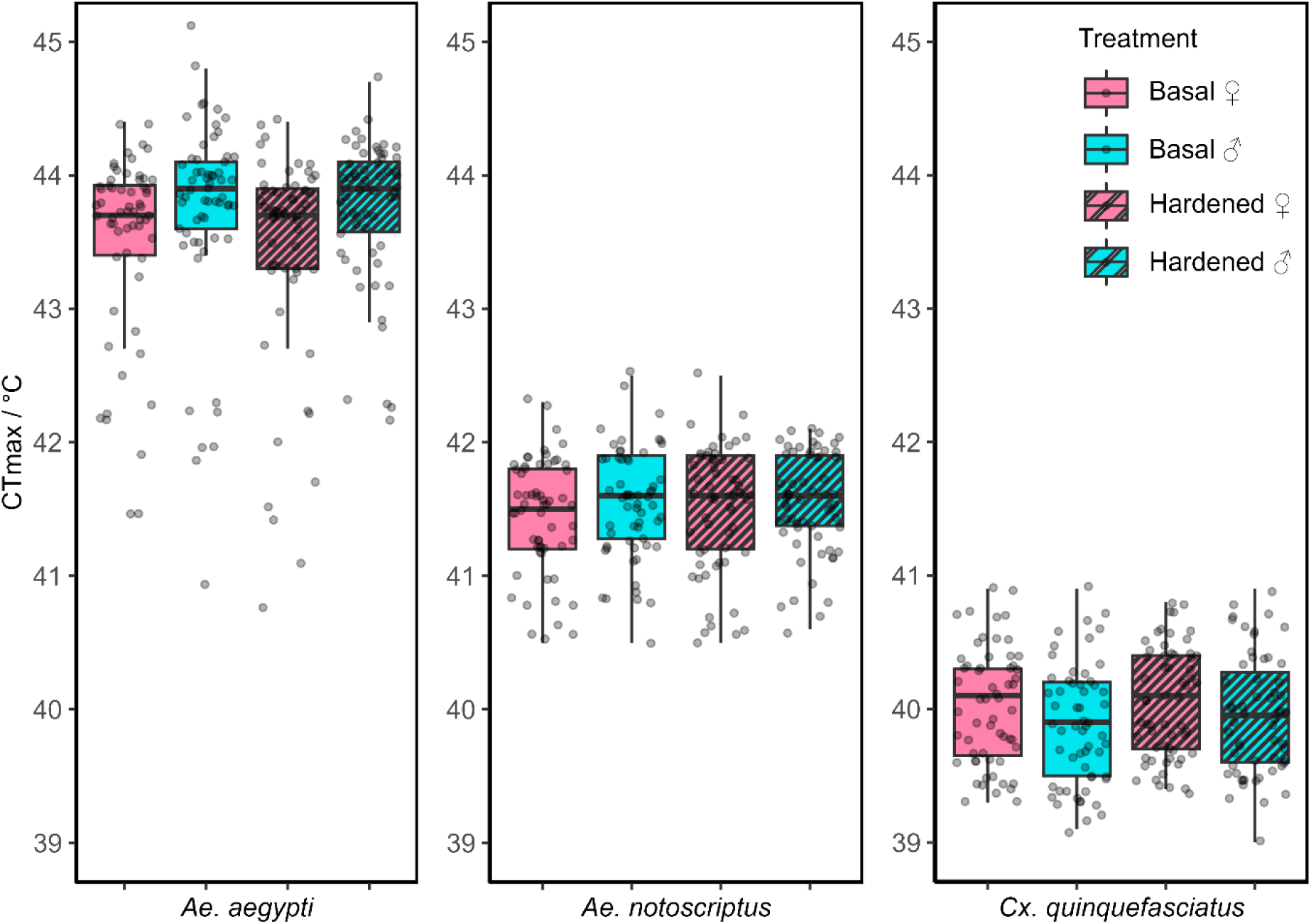
CTmax of basal and hardened male (♂) and female (♀) *Ae. aegypti*, *Ae. notoscriptus* and *Cx. quinquefasciatus*. CTmax was tested at a ramping rate of 0.1 °C per minute from an initial temperature of 26 °C. Grey overlaid data points represent individual CTmax measurements. Pink and blue boxplots indicate females and males respectively. Striped boxplots represent hardened mosquitoes, while plain boxplots represent basal mosquitoes.

### Heat hardening effects on heat knockdown time

The effect of heat hardening (one-hour heat shock at 41 °C) on heat knockdown time at a constant temperature near the upper thermal limit (CTmax) was assessed separately in male and female mosquitoes from the three species (Figure 4). Though heat hardening increased heat knockdown time by 0.5 minutes in *Ae. notoscriptus* and 0.6 minutes in *Cx. quinquefasciatus,* and it decreased by 1.0 minutes in *Ae. aegypti,* none of these effects were statistically significant (all *p*-values > 0.081) (Table S4). A sex difference in heat tolerance was observed in *Cx. quinquefasciatus,* with males showing reduced tolerance compared to females (Δ = - 1.895, *F*_1, 234_ = 13.415, *p* < 0.001). In contrast, differences between the sexes in *Ae. aegypti* (Δ = 0.777, *F*_1, 236_ = 1.992) and *Ae. notoscriptus* Δ = - 0.394, *F*_1, 236_ = 0.778) were not statistically significant (all *p*-values > 0.159; Figure S4). Similar to CTmax, mosquito age had a significant effect on heat tolerance in terms of heat knockdown time in all three species (*F*_2, 235_ = 4.519, *p* = 0.035 for *Ae. aegypti*; *F*_2, 235_ = 14.074, *p* < 0.001 for *Ae. notoscriptus*; *F*_3, 232_ = 55.846, *p* < 0.001 for *Cx. quinquefasciatus*). While *Ae. aegypti* and *Ae. notoscriptus* showed decreased heat tolerance with increasing age, *Cx. quinquefasciatus* exhibited increased heat tolerance in older individuals. Furthermore, two-way and three-way interaction effects among age, sex, and hardening treatment in each species did not significantly influence heat knockdown time (all *p*-values > 0.062; Table S4).

**Figure 4.**
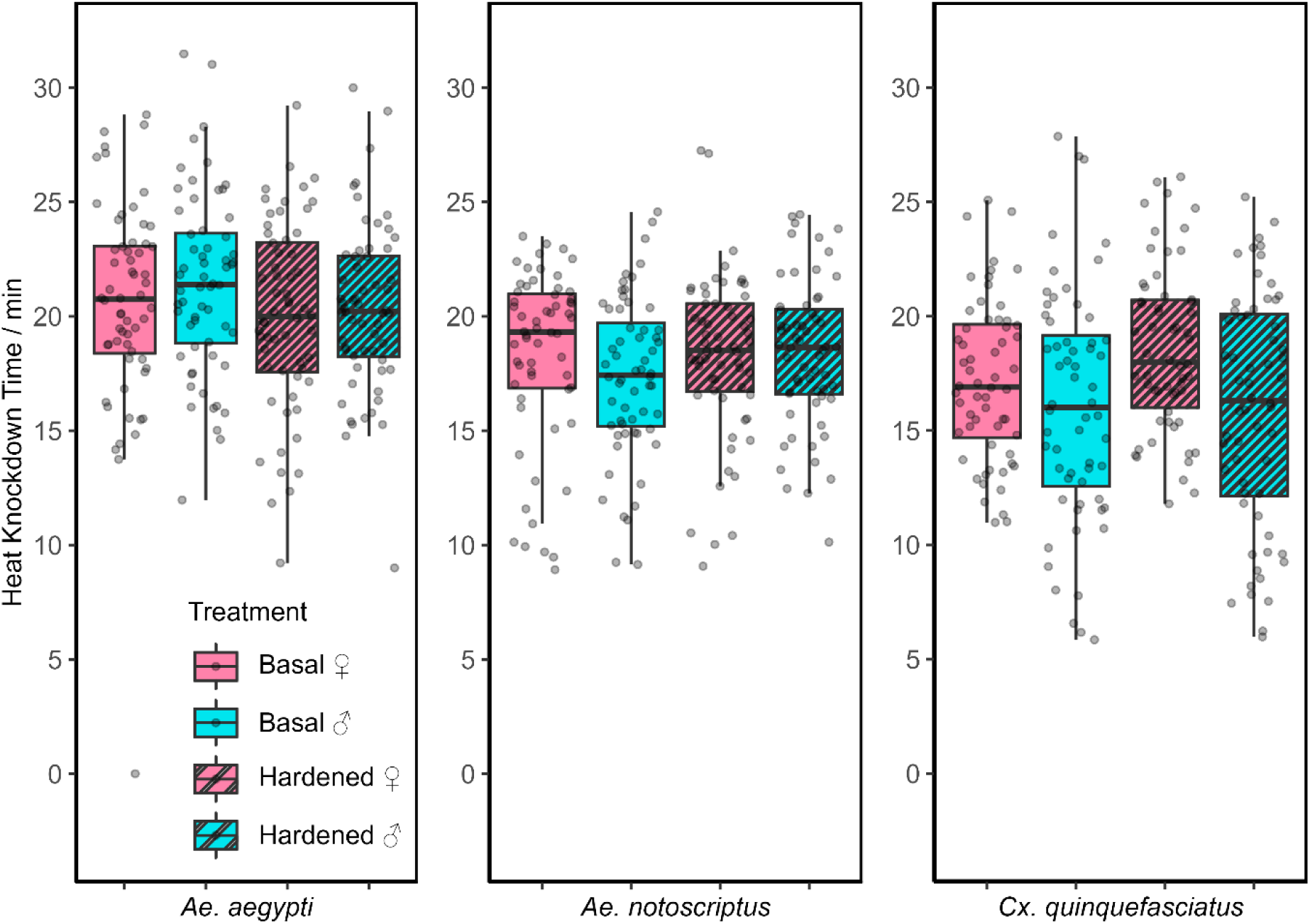
Heat knockdown time of basal and hardened male (♂) and female (♀) *Ae. aegypti*, *Ae. notoscriptus*, and *Cx. quinquefasciatus* mosquitoes. Heat knockdown time was measured at constant temperatures of 43.1 °C for *Ae. aegypti*, 41.2 °C for *Ae. notoscriptus*, and 39.8°C for *Cx. quinquefasciatus.* Grey overlaid data points represent individual measurements; Pink and blue boxplots indicate females and males respectively. Striped boxplots represent hardened mosquitoes, while plain boxplots represent basal mosquitoes.

### Cross-generational effects of heat shock exposure in *Aedes aegypti*

Cross-generational effects of heat exposure on male and female *Aedes aegypti* were assessed by comparing heat shock knockdown (Figure 5A), heat shock mortality (Figure 5B), CTmax (Figure 6A) and heat knockdown time (Figure 6B) in the offspring of female parents exposed to heat (one-hour heat shock at 41 °C). There were no significant differences in heat knockdown or mortality between the offspring of treated and untreated female parents for either sex or trait (all *p* values > 0.128; Table S5), with the exception of knockdown tolerance in male offspring, which was significantly reduced following female parental heat exposure (ΔLT_50_ = - 0.345 °C, t_18_ = 4.430, *p* < 0.001, Figure 5A; Tables 3 and S5). Female offspring had significantly higher heat tolerance than male offspring in both the heat knockdown and mortality assays (Figure 5, Table 3) regardless of female parental heat exposure condition (all *p* values < 0.002; Tables 3 and S5), except for knockdown tolerance in the offspring of untreated female parents (ΔLT_50_ = 0.136 °C, t_18_ = 1.839, *p* = 0.067; Tables 3 and S5).

**Figure 5.**
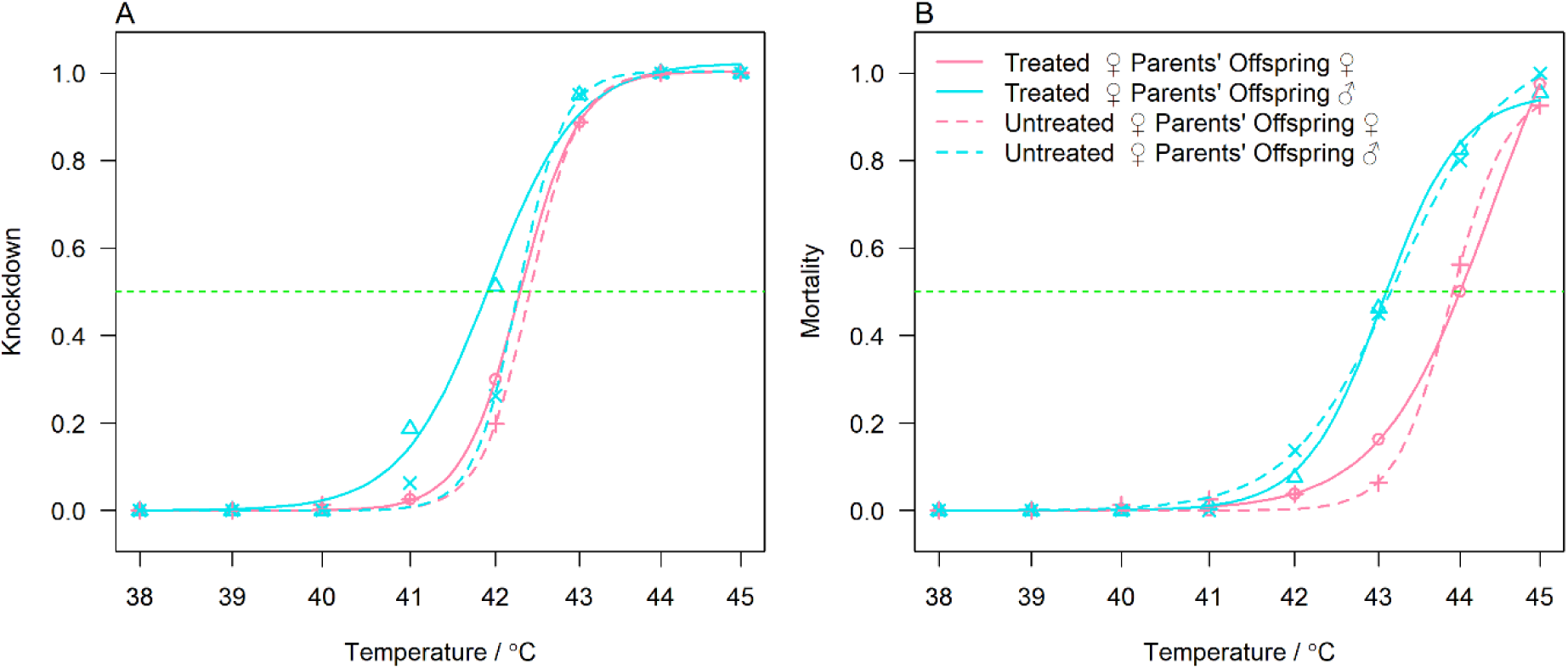
Cross-generational effects of heat shocks on heat knockdown (A) and 24-hr mortality (B) curves for male (♂) and female (♀) *Ae. aegypti*. Offspring were tested from heat-treated (one-hour heat shock at 41°C) and untreated mated female parents. Dots represent the mean percentage of knockdown or mortality averaged across 8 replicates per treatment, sex and temperature. Lines represent the predicted probability of knockdown or mortality across temperatures: pink lines for female offspring and blue lines for male offspring. Solid lines indicate offspring from treated parents and dashed lines indicate offspring from untreated parents. Intersections with green lines represent LT_50_s for knockdown and mortality.

**Figure 6.**
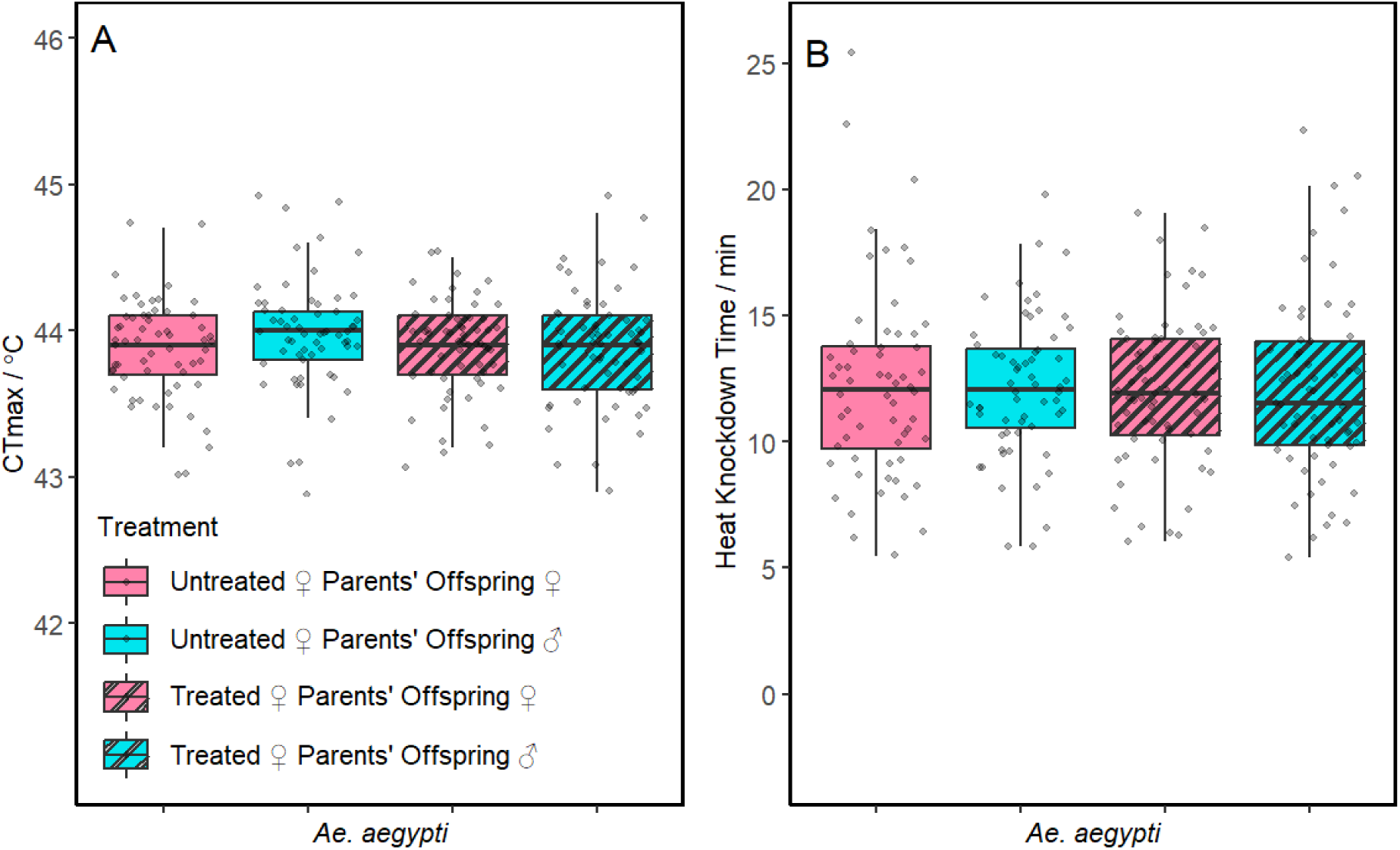
Cross-generational effects of heat shocks on CTmax (A) and heat knockdown time (B) of male (♂) and female (♀) *Ae. aegypti*. Offspring were tested from heat-treated (one-hour heat shock at 41 °C) and untreated female parents. Grey overlaid data points represent individual CTmax (A) and heat knockdown time (B) measurements. Pink indicates female offspring and blue indicates male offspring. Striped boxplots represent offspring from treated parents, while plain boxplots represent offspring from basal parents. In the CTmax assay, the initial temperature was 26 °C with a ramping rate of 0.1 °C per minute. In the heat knockdown time assay, mosquitoes were exposed to a constant temperature of 43.1 °C.

**Table 3:** Estimated LT_50_ (°C) values for cross-generational heat knockdown and 24-hour mortality in male and female *Ae. aegypti* offspring from heat-treated (one-hour heat shock at 41 °C) or untreated mated female parents.

|  | Female parental heat treatment | Offspring sex | LT<sub>50</sub> | Std. Error | Lower 95% CI | Upper 95% CI |
| --- | --- | --- | --- | --- | --- | --- |
| Knockdown |  |  |  |  |  |  |
|  | Untreated | Female | 42.406 | 0.052 | 42.304 | 42.508 |
|  | Untreated | Male | 42.270 | 0.052 | 42.167 | 42.373 |
|  | Treated | Female | 42.294 | 0.052 | 42.192 | 42.396 |
|  | Treated | Male | 41.925 | 0.057 | 41.812 | 42.038 |
| Mortality |  |  |  |  |  |  |
|  | Untreated | Female | 43.883 | 0.074 | 43.737 | 44.030 |
|  | Untreated | Male | 43.223 | 0.146 | 42.936 | 43.510 |
|  | Treated | Female | 44.392 | 0.410 | 43.583 | 45.200 |
|  | Treated | Male | 43.051 | 0.099 | 42.856 | 43.246 |

In contrast to the heat knockdown and mortality assay, CTmax remained consistent across offspring sex (*F*_1, 236_ = 1.411, *p* = 0.236) and female parental heat exposure conditions (*F*_1, 236_ = 8.017, *p* = 0.372; Figure 6A, Tables S6). Similarly, heat knockdown time was not significantly affected by female parental heat exposure (*F*_1, 237_ = 0.074, *p* = 0.786) or offspring sex (*F*_1, 237_ = 0.005, *p* = 0.944, Figure 6B; Tables S6). Additionally, there were no statistically significant interaction effects between offspring sex and female parental heat exposure for either CTmax or heat knockdown time (all *p* values > 0.183; Tables S6).

## Discussion

In this study, we evaluated the effect of heat hardening on the adult heat tolerance of three vector mosquito species - *Ae. aegypti*, *Ae. notoscriptus* and *Cx*. *quinquefasciatus* - by assessing multiple related traits. Our results revealed well-pronounced species differences, sexual dimorphism and evidence of age-related decline in heat tolerance, but acute heat hardening provided inconsistent benefits across traits. We also assessed cross-generational effects of female parental acute heat exposure and found a weak, sex-specific negative impact in *Ae. aegypti* offspring. Overall, our findings highlight the complexity of mosquito thermal biology, demonstrating substantial variation in heat tolerance across species, sexes, ages and thermal traits.

### Heat hardening offers limited, but trait-specific benefits

Our results demonstrate that the effects of heat hardening on heat tolerance are both species-specific and trait-specific, consistent with previous findings in other insects species (Kellermann & Sgrò, 2018; Kellett et al., 2005). Heat hardening did not increase CTmax or static heat knockdown time in any species, but it substantially improved the static knockdown temperature in most cases. This could provide short-term resilience against acute thermal stress, delaying immobilization and allowing mosquitoes sufficient time to escape extreme heat by moving into a cooler microhabitats (Sauer et al., 2021). *Aedes notoscriptus* was the only species to exhibit improved heat survival following heat hardening. Species-specific physiological mechanisms may influence thermal plasticity (Lahondere & Bonizzoni, 2022), as has also been reported in other insects such as in *Drosophila* species (Baleba et al., 2024; Dahlgaard et al., 1998). A commonly discussed mechanism involves the regulation of heat stress, which even during hardening, is primarily achieved through the inducible expression of heat shock protein (HSPs) (Lindquist, 1981; Sivan et al., 2017). In mosquitoes, the level of HSP70 expression is particularly high under thermal stress (Benoit et al., 2011; Gross et al., 2009). These proteins function as molecular chaperones, preserving protein integrity under thermal stress (Hu et al., 2022), and their expression levels vary across species, traits and sexes (Singh et al., 2025; Sivan et al., 2017).

The absence of thermal plasticity in static knockdown temperature in *Ae. aegypti* males, and the relatively small hardening effect in *Ae. aegypti* females, may reflect evolutionary adaptation in this species to consistently warm tropical environments. In such settings, selection pressures may have already shaped a high baseline heat tolerance (Stillman, 2003; van Heerwaarden & Kellermann, 2020), leaving limited scope for further enhancement through short-term hardening (Overgaard et al., 2011; van Heerwaarden et al., 2016; Van Heerwaarden et al., 2024). Because hardening effects are inherently transient and expected to diminish over time (Dahlgaard et al., 1998; Sørensen et al., 2019), the 24-hour interval we employed between hardening treatment and experimental assays in our study may have contributed to dilution or disappearance of any heat hardening effect. Differences in hardening responses across species could also reflect threshold shifts, where species with higher basal heat tolerance require more intense hardening treatments to induce the greatest benefits (Van Heerwaarden et al., 2024). Further research could consider factors such as hardening duration, intensity and recovery interval.

### Sexual dimorphism in heat tolerance is consistent across species but trait specific

We observed sexual dimorphism in thermal tolerance across species, particularly in mortality and knockdown traits, with females exhibiting greater heat tolerance. This pattern has previously been reported in *Ae. aegypti* (Andersen et al., 2006; Chura et al., 2023) and *Cx. quinquefasciatus* (Chura et al., 2023), as well as in other vector mosquito species such as *Anopheles arabiensis* (Lyons et al., 2012) and *An. funestus* (Lyons et al., 2016; Lyons et al., 2012). The sex difference be due to their comparatively larger body size (Baudier et al., 2015; Lyberger et al., 2025), which can reduce water loss and associated dehydration under heat stress (Cossins & Bowler, 2012). In addition, females may have evolved enhanced heat tolerance due to exposure to acute heat stress during blood feeding on warm-blooded hosts (Benoit & Denlinger, 2017; Lahondère & Lazzari, 2013). Male vulnerability to heat stress poses a concern under rising global temperatures, as sex-based vector control strategies including the sterile insect technique and incompatible *Wolbachia* release programs rely on male mosquito survival (Bouyer, 2024; Dobson, 2021; Ross et al., 2023). Therefore, detailed investigation of sex-specific differences in thermal tolerance in vector mosquitoes is essential for both ecological understanding and applied control programs.

The effects of sex and hardening on heat tolerance were not consistent across the different assays, with stronger effects in the heat knockdown and survival experiment. This discrepancy may be explained by differences in humidity among experimental setups. Specially, the incubators used for the heat knockdown and mortality experiments were relatively dry (RH ∼10% at 41°C), leading to combined heat and desiccation stress, whereas the vials employed in the CTmax and heat knockdown assays were set up at ambient humidity (RH ∼45%). It is also plausible that differences reflect the way traits were scored, given that knockdown occurred at temperatures 1-2°C lower than mortality, and cessation of movement in CTmax assays does not correlate well with mortality (Hoffmann et al., 1997; Ørsted et al., 2024).

Our findings also indicate that younger mosquitoes are more resilient to heat stress than older individuals, which is consistent with previous studies (Lyons et al., 2016; Lyons et al., 2012). However, we did not detect any interaction between age and sex. More controlled experiments that minimise block effects are recommended to confirm age related variation in thermal tolerance across species and to assess how it may be modulated by sex.

### Species-specific thermal tolerance reflects their ecological origin

Among the limited studies comparing species differences in thermal tolerance in mosquitoes, our findings reveal clear interspecific variation among *Ae. aegypti*, *Ae. notoscriptus* and *Cx*. *quinquefasciatus* across all four thermal traits tested, irrespective of sex or hardening condition. *Ae. aegypti* consistently exhibited the highest heat tolerance under both acute and ramped heat stress, followed by *Ae. notoscriptus*, with *Cx. quinquefasciatus* demonstrating the lowest tolerance. Similarly, Chura et al. (2023) reported higher heat tolerance in *Ae. aegypti* relative to *Cx*. *quinquefasciatus* based on LT_50_ in mortality endpoints. The discrepancies in absolute LT_50_ values between their study and ours may reflects differences in mortality assessment criteria. Chura et al. (2023) defined mortality as the absence of coordinated movement such as flying or climbing, whereas our study employed a stricter definition of complete immobility without recovery. Despite these methodological differences, both studies consistently indicate an approximately 3°C thermal tolerance gap between two species. This species level difference in thermal tolerance suggests that, *Ae. aegypti* may outperform *Cx*. *quinquefasciatus* and *Ae. notoscriptus* in regions where environmental temperature reaches extreme levels.

CTmax estimates in our study for *Ae. aegypti* (44.3 °C) were comparable to the 43. 9 °C reported by Pekľanská et al. (2025), though higher than the 40 °C reported by Bar-Zeev (1957). Beyond adults, previous work has also demonstrated superior thermal tolerance in *Ae. aegypti* larvae compared to certain *Anopheles* and *Culex* species (Buxton et al., 2020). This elevated thermal tolerance in *Ae. aegypti* may reflects its tropical origin (Deutsch et al., 2008; Hoffmann et al., 2013; Lahondere & Bonizzoni, 2022) while the lower thermal tolerance observed in *Ae. notoscriptus* may reflect its temperate origin (Duchemin et al., 2017; Metzger et al., 2022). Few studies have examined the heat tolerance of *Ae. notoscriptus* (Williams & Rau, 2011), with our study being the first to estimate adult CTmax, and the first comparative study assessing adult thermal tolerance across multiple traits. In contrast to this comparison, the low thermal tolerance observed in *Cx. quinquefasciatus* is unexpected, given its likely tropical-subtropical origin (Negi & Verma, 2018), which might typically suggest an intermediate level of thermal tolerance. However, differences in upper thermal limits of insect species may be minor unless extremely high latitudes are considered in the comparison (Hoffmann et al 2013), although this may not apply at the population level (Vorhees et al., 2013), and would be worth exploring further, particularly in *Cx. quinquefasciatus* and *Ae. notoscriptus*.

Most existing mosquito thermal tolerance studies have relied on CTmax as the primary endpoint (Bar-Zeev, 1957; Buxton et al., 2020; Pekľanská et al., 2025), whereas mortality-based LT_50_ values have been used less frequently (Chura et al., 2023; Sivan et al., 2020). To our knowledge, our study is the first to incorporate knockdown-based LT_50_ endpoints. The underutilization of knockdown-based LT_50_ endpoints may reflect the subjectivity in defining knockdown thresholds, whereas CTmax provides a more standardised and reproducible measure across studies. Moreover, the differences we observed between mortality and knockdown based LT_50_ values suggest that knockdown may underestimate actual heat tolerance.

### Cross-generation effects are weak and sex dependent

Parental heat shock exposure appeared to have only minimal cross-generational effects on *Ae. aegypti* offspring thermal tolerance, with the sole detectable impact being reduced static knockdown tolerance in male offspring. This suggests a potential negative carryover effect that imposes physiological costs on subsequent generations (Foo et al., 2019; Mpofu et al., 2022; Sales et al., 2021). Although we did not detect any cross-generational plastic effect on CTmax or static heat knockdown time in mosquitoes, adaptive plastic responses have been identified in other species including *Drosophila mojavensis* (Diaz et al., 2021), highlighting the possibility of species-specificity in cross-generational plasticity. Currently, evidence for cross-generational effect of parental heat stress in vector mosquitoes is limited. While some studies have identified beneficial cross-generational effects of heat stress (Dennington et al., 2024; Pekľanská et al., 2025), these were not designed to separate plastic effects from those of evolutionary changes due to direct selection on heat tolerance.

## Conclusion

Our study emphasizes the need to carefully consider context-specific patterns of heat hardening and sexual dimorphism in thermal tolerance when predicting the impact of heat stress on mosquitoes. Incorporating species, sex and context specific thermal responses into models of mosquito distribution and fitness under rising temperature is essential (Couper et al., 2024), as mosquito overall fitness (Bagni et al., 2024; Ciota et al., 2014) and vector biology (Onyango et al., 2020; Paaijmans et al., 2012) will be affected by environmental temperature. The hardening effects we have investigated here may also be relevant to implementing mosquito control strategies such as those based on releasing males that generate sterility in field females and need to be competitive with field males already present in a population (Ross et al., 2019).

## Acknowledgements

We thank Leon Hugo and Scott Ritchie for providing mosquito populations used in the study. We also thank Miriama Pekľanská, Ella Yeatman and Mel Berran for providing technical assistance with experiments.

## Author contributions

Conceptualization: A.L.W., B.v.H., A.A.H., P.A.R.; Formal analysis: A.L.W, B.v.H., A.A.H., P.A.R.; Funding acquisition: B.v.H., A.A.H., P.A.R.; Investigation: A.L.W.; Methodology: A.L.W., B.v.H., P.A.R.; Supervision: B.v.H., A.A.H., P.A.R.; Visualization: A.L.W; Writing – original draft: A.L.W., B.v.H., A.A.H., P.A.R.; Writing – review & editing: A.L.W., B.v.H., A.A.H., P.A.R.

## References

AGDH. (2025). Australian Government Department of Health and Aged Care, mosqito-born diseases. Retrieved July, 15^th^ from https://www.health.gov.au/node/44804?query=mosqito-born+diseases&search_scope=0

Andersen, J. P., Schwartz, A., Gramsbergen, J. B., & Loeschcke, V. (2006). Dopamine levels in the mosquito *Aedes aegypti* during adult development, following blood feeding and in response to heat stress. J Insect Physiol, 52(11-12), 1163–1170. https://www.sciencedirect.com/science/article/abs/pii/S0022191006001545?via%3Dih <u>ub</u>

Arnold, P. A., Nicotra, A. B., & Kruuk, L. E. B. (2019). Sparse evidence for selection on phenotypic plasticity in response to temperature. Philos Trans R Soc Lond B Biol Sci, 374(1768), 20180185. 10.1098/rstb.2018.0185

Auguie, B., & Antonov, A. (2017). gridExtra: Miscellaneous functions for" Grid" Graphics. (R package version 2.3) [Computer software]. In (pp. 1–9).

Bagni, T., Siaussat, D., Maria, A., Fuentes, A., Couzi, P., & Massot, M. (2024). Fitness under high temperatures is overestimated when daily thermal fluctuation is ignored. J Therm Biol, 103806.

Baleba, S. B., Jiang, N.-J., & Hansson, B. S. (2024). Temperature-mediated dynamics: Unravelling the impact of temperature on cuticular hydrocarbon profiles, mating behaviour, and life history traits in three *Drosophila* species. Heliyon, 10(17).

Bar-Zeev, M. (1957). The effect of extreme temperatures on different stages of *Aedes aegypti* (L.). Bull Entomol Res, 48(3), 593–599.

Barr, J. S., Martin, L. E., Tate, A. T., & Hillyer, J. F. (2024). Warmer environmental temperature accelerates aging in mosquitoes, decreasing longevity and worsening infection outcomes. Immun Ageing, 21(1), 61. 10.1186/s12979-024-00465-w

Bates, D., Mächler, M., Bolker, B., & Walker, S. (2015). Fitting linear mixed-effects models using lme4 (R package version 1.1.35.3) [Computer software]. J Stat Softw, 67, 1–48. https://cran.r-project.org/package=lme4

Baudier, K. M., Mudd, A. E., Erickson, S. C., & O’Donnell, S. (2015). Microhabitat and body size effects on heat tolerance: implications for responses to climate change (army ants: Formicidae, Ecitoninae). J Anim Ecol, 84(5), 1322–1330.

Benoit, J. B., & Denlinger, D. L. (2017). Bugs battle stress from hot blood. Elife, 6, e33035. https://pmc.ncbi.nlm.nih.gov/articles/PMC5697928/

Benoit, J. B., Lopez-Martinez, G., Patrick, K. R., Phillips, Z. P., Krause, T. B., & Denlinger, D. L. (2011). Drinking a hot blood meal elicits a protective heat shock response in mosquitoes. Proc Natl Acad Sci U S A, 108(19), 8026–8029. 10.1073/pnas.1105195108

Bouyer, J. (2024). Current status of the sterile insect technique for the suppression of mosquito populations on a global scale. Infect Dis Poverty, 13(1), 68. https://idpjournal.biomedcentral.com/counter/pdf/10.1186/s40249-024-01242-z.pdf

Bowler, K. (2005). Acclimation, heat shock and hardening. J Therm Biol, 30(2), 125–130.

Brady, O. J., & Hay, S. I. (2020). The global expansion of dengue: How *Aedes aegypti* mosquitoes enabled the first pandemic arbovirus. Annu Rev Entomol, 65, 191–208. 10.1146/annurev-ento-011019-024918

Brady, O. J., Johansson, M. A., Guerra, C. A., Bhatt, S., Golding, N., Pigott, D. M., Delatte, H., Grech, M. G., Leisnham, P. T., & Maciel-de-Freitas, R. (2013). Modelling adult *Aedes aegypti* and *Aedes albopictus* survival at different temperatures in laboratory and field settings. Parasit Vectors, 6, 1–12.

Bujan, J., Roeder, K. A., Yanoviak, S. P., & Kaspari, M. (2020). Seasonal plasticity of thermal tolerance in ants. Ecol, 101(6), e03051.

Buxton, M., Nyamukondiwa, C., Dalu, T., Cuthbert, R. N., & Wasserman, R. J. (2020). Implications of increasing temperature stress for predatory biocontrol of vector mosquitoes. Parasit Vectors, 13(1), 604.

Carlassara, M., Khorramnejad, A., Oker, H., Bahrami, R., Lozada-Chávez, A. N., Mancini, M. V., Quaranta, S., Body, M. J. A., Lahondère, C., & Bonizzoni, M. (2024). Population-specific responses to developmental temperature in the arboviral vector *Aedes albopictus*: Implications for climate change. Glob Chang Biol, 30(3), e17226. 10.1111/gcb.17226

Carrington., L. B., Armijos, M. V., Lambrechts, L., Barker, C. M., & Scott, T. W. (2013). Effects of fluctuating daily temperatures at critical thermal extremes on *Aedes aegypti* life-history traits. PLoS One, 8(3), e58824. 10.1371/journal.pone.0058824

Chura, M., Healy, K., Diaz, R., & Kaller, M. (2023). Effects of species, sex, and diet on thermal tolerance of *Aedes aegypti* and *Culex quinquefasciatus* (Diptera: Culicidae). J Med Entomol, 60(4), 637–643. 10.1093/jme/tjad037

Ciota, A. T., Matacchiero, A. C., Kilpatrick, A. M., & Kramer, L. D. (2014). The effect of temperature on life history traits of *Culex* mosquitoes. J Med Entomol, 51(1), 55–62. 10.1603/me13003

Claflin, S. B., & Webb, C. E. (2015). Ross River virus: Many vectors and unusual hosts make for an unpredictable pathogen. PLoS Pathog, 11(9), e1005070. 10.1371/journal.ppat.1005070

Cointe, B., & Guillemot, H. (2023). A history of the 1.5 C target. Wiley Interdiscip Rev Clim Change, 14(3), e824.

Colinet, H., Sinclair, B. J., Vernon, P., & Renault, D. (2015). Insects in fluctuating thermal environments. Annu Rev Entomol, 60, 123–140. 10.1146/annurev-ento-010814-021017

Cossins, A. R., & Bowler, K. (2012). Temperature biology of animals. Chapman and Hall.

Couper, L., Farner, J., Caldwell, J., Childs, M., Harris, M., Kirk, D., Nova, N., Shocket, M., Skinner, E., Uricchio, L., Exposito-Alonso, M., & Mordecai, E. (2021). How will mosquitoes adapt to climate warming? Elife, 10, e69630.

Couper, L. I., Dodge, T. O., Hemker, J. A., Kim, B. Y., Exposito-Alonso, M., Brem, R. B., Mordecai, E. A., & Bitter, M. C. (2024). Evolutionary adaptation under climate change: *Aedes* sp. demonstrates potential to adapt to warming. bioRxiv. 10.1101/2024.08.23.609454

Dahlgaard, J., Loeschcke, V., Michalak, P., & Justesen, J. (1998). Induced thermotolerance and associated expression of the heat-shock protein Hsp70 in adult *Drosophila melanogaster*. Funct Ecol, 12(5), 786–793.

de Souza, W. M., & Weaver, S. C. (2024). Effects of climate change and human activities on vector-borne diseases. Nat Rev Microbiol, 22(8), 476–491. 10.1038/s41579-024-01026-0

Delatte, H., Gimonneau, G., Triboire, A., & Fontenille, D. (2009). Influence of temperature on immature development, survival, longevity, fecundity, and gonotrophic cycles of *Aedes albopictus*, vector of chikungunya and dengue in the Indian Ocean. J Med Entomol, 46(1), 33–41. 10.1603/033.046.0105

Dennington, N. L., Grossman, M. K., Ware-Gilmore, F., Teeple, J. L., Johnson, L. R., Shocket, M. S., McGraw, E. A., & Thomas, M. B. (2024). Phenotypic adaptation to temperature in the mosquito vector, *Aedes aegypti*. Glob Chang Biol, 30(1), e17041. 10.1111/gcb.17041

Deutsch, C. A., Tewksbury, J. J., Huey, R. B., Sheldon, K. S., Ghalambor, C. K., Haak, D. C., & Martin, P. R. (2008). Impacts of climate warming on terrestrial ectotherms across latitude. Proc Natl Acad Sci USA, 105(18), 6668–6672.

Diaz, F., Kuijper, B., Hoyle, R. B., Talamantes, N., Coleman, J. M., & Matzkin, L. M. (2021). Environmental predictability drives adaptive within-and transgenerational plasticity of heat tolerance across life stages and climatic regions. Funct Ecol, 35(1), 153–166.

Dobson, S. L. (2021). When more is less: Mosquito population suppression using sterile, incompatible and genetically modified male mosquitoes. J Med Entomol, 58(5), 1980–1986.

Duchemin, J. B., Mee, P. T., Lynch, S. E., Vedururu, R., Trinidad, L., & Paradkar, P. (2017). Zika vector transmission risk in temperate Australia: a vector competence study. Virol J, 14(1), 108. 10.1186/s12985-017-0772-y

Ezeakacha, N. F., & Yee, D. A. (2019). The role of temperature in affecting carry-over effects and larval competition in the globally invasive mosquito *Aedes albopictus*. Parasit Vectors, 12, 1–11.

Ferguson, L. F., Ross, P. A., & van Heerwaarden, B. (2024). *Wolbachia* infection negatively impacts *Drosophila simulans* heat tolerance in a strain-and trait-specific manner. Environ Microbiol, 26(4), e16609.

Foo, I. J., Hoffmann, A. A., & Ross, P. A. (2019). Cross-generational effects of heat stress on fitness and *Wolbachia* density in *Aedes aegypti* mosquitoes. Trop Med Infect Dis, 4(1). 10.3390/tropicalmed4010013

Fox, C. W., & Mousseau, T. A. (1998). Maternal effects as adaptations for transgenerational phenotypic plasticity in insects. In C. W. Fox & T. A. Mousseau (Eds.), Maternal effects as adaptations (Vol. 159, pp. 159–177). Oxford Academic.

Fox, J., & Weisberg, S. (2023). car: companion to applied regression.(R package version 3.1.2) [Computer software]. https://cran.r-project.org/package=car, 2023. https://cran.r-project.org/web/packages/car/index.html

Goindin, D., Delannay, C., Ramdini, C., Gustave, J., & Fouque, F. (2015). Parity and longevity of *Aedes aegypti* according to temperatures in controlled conditions and consequences on dengue transmission risks. PLoS One, 10(8), e0135489. 10.1371/journal.pone.0135489

Gray, E. M. (2013). Thermal acclimation in a complex life cycle: the effects of larval and adult thermal conditions on metabolic rate and heat resistance in *Culex pipiens* (Diptera: Culicidae). J Insect Physiol, 59(10), 1001–1007. 10.1016/j.jinsphys.2013.08.001

Grimm, N. B., Chapin III, F. S., Bierwagen, B., Gonzalez, P., Groffman, P. M., Luo, Y., Melton, F., Nadelhoffer, K., Pairis, A., & Raymond, P. A. (2013). The impacts of climate change on ecosystem structure and function. Front Ecol Environ, 11(9), 474–482.

Gross, T. L., Myles, K. M., & Adelman, Z. N. (2009). Identification and characterization of heat shock 70 genes in Aedes aegypti (Diptera: Culicidae). J Med Entomol, 46(3), 496–504. 10.1603/033.046.0313

Gunderson, A. R., Dillon, M. E., Stillman, J. H., & Trullas, S. C. (2017). Estimating the benefits of plasticity in ectotherm heat tolerance under natural thermal variability. Funct Ecol, 31(8), 1529–1539. 10.1111/1365-2435.12874

Harrell, F. E., & Dupont, C. (2023). Hmisc: Harrell miscellaneous. (R package version 5.1-2) [Computer software]. R Found. Stat. Comput. https://cran.r-project.org/web/packages/Hmisc/index.html

Harvey, J. A., Heinen, R., Gols, R., & Thakur, M. P. (2020). Climate change-mediated temperature extremes and insects: From outbreaks to breakdowns. Glob Chang Biol, 26(12), 6685–6701. https://pmc.ncbi.nlm.nih.gov/articles/PMC7756417/

Hoffmann, A. A., & Bridle, J. (2022). The dangers of irreversibility in an age of increased uncertainty: revisiting plasticity in invertebrates. Oikos, 2022(4), e08715.

Hoffmann, A. A., Chown, S. L., & Clusella-Trullas, S. (2013). Upper thermal limits in terrestrial ectotherms: how constrained are they? Funct Ecol, 27(4), 934–949.

Hoffmann, A. A., Dagher, H., Hercus, M., & Berrigan, D. (1997). Comparing different measures of heat resistance in selected lines of *Drosophila melanogaster*. J Insect Physiol, 43(4), 393–405.

Hoffmann, A. A., Sgrò, C. M., & van Heerwaarden, B. (2023). Testing evolutionary adaptation potential under climate change in invertebrates (mostly *Drosophila*): findings, limitations and directions. J Exp Biol, 226(14), jeb245749.

Hoffmann, A. A., Sørensen, J. G., & Loeschcke, V. (2003). Adaptation of *Drosophila* to temperature extremes: bringing together quantitative and molecular approaches. J Therm Biol, 28(3), 175–216.

Hu, C., Yang, J., Qi, Z., Wu, H., Wang, B., Zou, F., Mei, H., Liu, J., Wang, W., & Liu, Q. (2022). Heat shock proteins: Biological functions, pathological roles, and therapeutic opportunities. Med Comm, 3(3), e161. https://pmc.ncbi.nlm.nih.gov/articles/PMC9345296/

Jansen, C. C., & Beebe, N. W. (2010). The dengue vector *Aedes aegypti*: what comes next. Microbes Infect, 12(4), 272–279. 10.1016/j.micinf.2009.12.011

Jorgensen, L. B., Malte, H., Orsted, M., Klahn, N. A., & Overgaard, J. (2021). A unifying model to estimate thermal tolerance limits in ectotherms across static, dynamic and fluctuating exposures to thermal stress. Sci Rep, 11(1), 12840. 10.1038/s41598-021-92004-6

Jørgensen, L. B., Malte, H., Overgaard, J., & White, C. (2019). How to assess *Drosophila* heat tolerance: Unifying static and dynamic tolerance assays to predict heat distribution limits. Funct Ecol, 33(4), 629–642. 10.1111/1365-2435.13279

Kamiya, T., Greischar, M. A., Wadhawan, K., Gilbert, B., Paaijmans, K., & Mideo, N. (2020). Temperature-dependent variation in the extrinsic incubation period elevates the risk of vector-borne disease emergence. Epidemics, 30. 10.1016/j.epidem.2019.100382

Kellermann, V., & Sgrò, C. M. (2018). Evidence for lower plasticity in CTMAX at warmer developmental temperatures. J Evol Biol, 31(9), 1300–1312.

Kellermann, V., & van Heerwaarden, B. (2019). Terrestrial insects and climate change: adaptive responses in key traits. Physiol Entomol, 44(2), 99–115.

Kellett, M., Hoffmann, A. A., & Mckechnie, S. W. (2005). Hardening capacity in the *Drosophila melanogaster* species group is constrained by basal thermotolerance. J Funct Ecol, 853-858.

Lahondere, C., & Bonizzoni, M. (2022). Thermal biology of invasive *Aedes* mosquitoes in the context of climate change. Curr Opin Insect Sci, 51, 100920. 10.1016/j.cois.2022.100920

Lahondère, C., & Lazzari, C. R. (2013). Thermal Stress and Thermoregulation During Feeding in Mosquitoes. In Anopheles mosquitoes - New insights into malaria vectors. 10.5772/56288

Lalejini, A., Ferguson, A. J., Grant, N. A., & Ofria, C. (2021). Adaptive phenotypic plasticity stabilizes evolution in fluctuating environments. Front Ecol Evol, 9. 10.3389/fevo.2021.715381

Lindquist, S. (1981). Regulation of protein synthesis during heat shock. Nature, 293(5830), 311–314. https://www.nature.com/articles/293311a0

Lyberger, K., Farner, J. E., Couper, L., & Mordecai, E. A. (2025). Plasticity in mosquito size and thermal tolerance across a latitudinal climate gradient. J Anim Ecol, 94(3), 330–339.

Lyons, C., Oliver, S., Hunt, R., & Coetzee, M. (2016). the influence of insecticide resistance, age, sex, and blood feeding frequency on thermal tolerance of wild and laboratory phenotypes of *Anopheles funestus* (Diptera: Culicidae). J Med Entomol, 53(2), 394–400.

Lyons, C. L., Coetzee, M., Terblanche, J. S., & Chown, S. L. (2012). Thermal limits of wild and laboratory strains of two African malaria vector species, *Anopheles arabiensis* and *Anopheles funestus*. Malar J, 11, 1–14.

Ma, C.-S., Ma, G., & Pincebourde, S. (2021). Survive a warming climate: insect responses to extreme high temperatures. Ann Rev Entomol, 66(1), 163–184.

MacLean, H. J., Sorensen, J. G., Kristensen, T. N., Loeschcke, V., Beedholm, K., Kellermann, V., & Overgaard, J. (2019). Evolution and plasticity of thermal performance: an analysis of variation in thermal tolerance and fitness in 22 *Drosophila* species. Philos Trans R Soc Lond B Biol Sci, 374(1778), 20180548. 10.1098/rstb.2018.0548

Mee, P. T., Buultjens, A. H., Oliver, J., Brown, K., Crowder, J. C., Porter, J. L., Hobbs, E. C., Judd, L. M., Taiaroa, G., & Puttharak, N. (2024). Mosquitoes provide a transmission route between possums and humans for Buruli ulcer in southeastern Australia. Nat Microbiol, 9(2), 377–389. https://www.nature.com/articles/s41564-023-01553-1.pdf

Metzger, M. E., Wekesa, J. W., Kluh, S., Fujioka, K. K., Saviskas, R., Arugay, A., McConnell, N., Nguyen, K., Krueger, L., Hacker, G. M., Hu, R., & Kramer, V. L. (2022). Detection and Establishment of *Aedes notoscriptus* (Diptera: Culicidae) Mosquitoes in Southern California, United States. J Med Entomol, 59(1), 67–77. 10.1093/jme/tjab165

Mike, F., Davis, T. L., & Wickham, H. (2023). ggpattern:‘ggplot2’pattern geoms (R package version 1.0.1) [Computer software]. In.

Mordecai, E. A., Caldwell, J. M., Grossman, M. K., Lippi, C. A., Johnson, L. R., Neira, M., Rohr, J. R., Ryan, S. J., Savage, V., Shocket, M. S., Sippy, R., Stewart Ibarra, A. M., Thomas, M. B., & Villena, O. (2019). Thermal biology of mosquito-borne disease. Ecol Lett, 22(10), 1690–1708. 10.1111/ele.13335

Mpofu, P., Machekano, H., & Nyamukondiwa, C. (2022). Parental acclimation reduces offspring thermal fitness in the postharvest insect species *Sitotroga cerealella* (Olivier). IOBC-WPRS Bulletin, 159, 53–58.

Muturi, E. J., Muriu, S., Shililu, J., Mwangangi, J. M., Jacob, B. G., Mbogo, C., Githure, J., & Novak, R. J. (2008). Blood-feeding patterns of *Culex quinquefasciatus* and other culicines and implications for disease transmission in Mwea rice scheme, Kenya. Parasitol Res, 102(6), 1329–1335. 10.1007/s00436-008-0914-7

Negi, C., & Verma, P. (2018). Review on *Culex quinquefasciatus*: Southern House Mosquito. Int J Life-sci Res, 4(1), 1563–1566. 10.21276/ijlssr.2018.4.1.9

Oliveira, B. F., Yogo, W. I., Hahn, D. A., Yongxing, J., & Scheffers, B. R. (2021). Community-wide seasonal shifts in thermal tolerances of mosquitoes. Ecol, 102(7), e03368.

Onyango, M. G., Bialosuknia, S. M., Payne, A. F., Mathias, N., Kuo, L., Vigneron, A., DeGennaro, M., Ciota, A. T., & Kramer, L. D. (2020). Increased temperatures reduce the vectorial capacity of *Aedes* mosquitoes for Zika virus. Emerg Microbes Infect, 9(1), 67–77. 10.1080/22221751.2019.1707125

Ørsted, M., Willot, Q., Olsen, A. K., Kongsgaard, V., & Overgaard, J. (2024). Thermal limits of survival and reproduction depend on stress duration: A case study of *Drosophila suzukii*. Ecol Lett, 27(3), e14421.

Overgaard, J., Kristensen, T. N., Mitchell, K. A., & Hoffmann, A. A. (2011). Thermal tolerance in widespread and tropical *Drosophila* species: does phenotypic plasticity increase with latitude? Am Nat, 178(S1), S80–S96.

Overgaard, J., Kristensen, T. N., & Sørensen, J. G. (2012). Validity of thermal ramping assays used to assess thermal tolerance in arthropods. PLoS One, 7(3). 10.1371/journal.pone.0032758

Paaijmans, K. P., Blanford, S., Chan, B. H., & Thomas, M. B. (2012). Warmer temperatures reduce the vectorial capacity of malaria mosquitoes. Biol Lett, 8(3), 465–468. 10.1098/rsbl.2011.1075

Pekľanská, M., van Heerwaarden, B., Hoffmann, A. A., Nouzová, M., Šíma, R., & Ross, P. A. (2025). Elevated developmental temperatures below the lethal limit reduce *Aedes aegypti* fertility. J Exp Biol, 228(3), JEB249803.

Reeves, W. C., Hardy, J. L., Reisen, W. K., & Milby, M. M. (1994). Potential effect of global warming on mosquito-borne arboviruses. J Med Entomol, 31(3), 323–332.

Ritz, C., Baty, F., Streibig, J. C., & Gerhard, D. (2015). Dose-response analysis using R (R package version 3.0.1). PLoS One, 10(12), e0146021. https://journals.plos.org/plosone/article/file?id=10.1371/journal.pone.0146021&type=p rintable

Rodrigues, Y. K., & Beldade, P. (2020). Thermal plasticity in insects’ response to climate change and to multifactorial environments. Front Ecol Evol, 8. 10.3389/fevo.2020.00271

Ross, P., Axford, J. K., Richardson, K. M., Endersby-Harshman, N. M., & Hoffmann, A. A. (2017). Maintaining *Aedes aegypti* mosquitoes Infected with *Wolbachia*. J Vis Exp(126). 10.3791/56124

Ross, P. A., Elfekih, S., Collier, S., Klein, M. J., Lee, S. S., Dunn, M., Jackson, S., Zhang, Y., Axford, J. K., & Gu, X. (2023). Developing *Wolbachia*-based disease interventions for an extreme environment. PLoS Pathog, 19(1), e1011117. https://journals.plos.org/plospathogens/article/file?id=10.1371/journal.ppat.1011117&type=printable

Ross, P. A., Turelli, M., & Hoffmann, A. A. (2019). Evolutionary ecology of *Wolbachia* releases for disease control. Annu Rev Genet, 53(1), 93–116.

RStudio Team. (2024). RStudio: Integrated development environment for R (Version 2024.04.1+748) [Computer software]. In *(No Title)*: Posit, PBC.

Sales, K., Vasudeva, R., & Gage, M. J. G. (2021). Fertility and mortality impacts of thermal stress from experimental heatwaves on different life stages and their recovery in a model insect. R Soc Open Sci, 8(3), 201717. 10.1098/rsos.201717

Sauer, F., Grave, J., Lühken, R., & Kiel, E. (2021). Habitat and microclimate affect the resting site selection of mosquitoes. Med Vet Entomol, 35(3), 379–388.

Scheffers, B. R., De Meester, L., Bridge, T. C., Hoffmann, A. A., Pandolfi, J. M., Corlett, R. T., Butchart, S. H., Pearce-Kelly, P., Kovacs, K. M., & Dudgeon, D. (2016). The broad footprint of climate change from genes to biomes to people. Science, 354(6313), aaf7671.

Sejerkilde, M., Sørensen, J. G., & Loeschcke, V. (2003). Effects of cold-and heat hardening on thermal resistance in *Drosophila melanogaster*. J Insect Physiol, 49(8), 719–726. https://www.sciencedirect.com/science/article/abs/pii/S0022191003000957?via%3Dih ub

Sgro, C. M., Terblanche, J. S., & Hoffmann, A. A. (2016). What can plasticity contribute to Insect responses to climate change? Annu Rev Entomol, 61, 433–451. 10.1146/annurev-ento-010715-023859

Singh, P., Pasi, S., Pande, V., & Dhiman, R. C. (2025). Heat Shock Proteins expression in malaria and dengue vector. Int J Biometeorol, 69(1), 225–232. https://link.springer.com/article/10.1007/s00484-024-02806-2

Sivan, A., Shriram, A. N., Muruganandam, N., & Thamizhmani, R. (2017). Expression of heat shock proteins (HSPs) in *Aedes aegypti* (L) and *Aedes albopictus* (Skuse) (Diptera: Culicidae) larvae in response to thermal stress. Acta Trop, 167, 121–127. 10.1016/j.actatropica.2016.12.017

Sivan, A., Shriram, A. N., Vanamail, P., & Sugunan, A. P. (2020). Thermotolerance and acclimation in the immature stages of *Aedes aegypti* (L) (Diptera: Culicidae) to simulated thermal stress. Int J Trop Insect Sci, 41(1), 333–344. 10.1007/s42690-020-00211-x

Sørensen, J. G., Kristensen, T. N., & Overgaard, J. (2016). Evolutionary and ecological patterns of thermal acclimation capacity in *Drosophila*: is it important for keeping up with climate change? Curr Opin Insect Sci, 17, 98–104. https://www.sciencedirect.com/science/article/abs/pii/S2214574516300967?via%3Dihub

Sørensen, M. H., Kristensen, T. N., Lauritzen, J. M. S., Noer, N. K., Høye, T. T., & Bahrndorff, S. (2019). Rapid induction of the heat hardening response in an Arctic insect. Biol Lett, 15(10), 20190613. https://pmc.ncbi.nlm.nih.gov/articles/PMC6832182/

Stillman, J. H. (2003). Acclimation capacity underlies susceptibility to climate change. Science, 301(5629), 65–65.

Urbanski, J. M., Benoit, J. B., Michaud, M. R., Denlinger, D. L., & Armbruster, P. (2010). The molecular physiology of increased egg desiccation resistance during diapause in the invasive mosquito, *Aedes albopictus*. Proc Biol Sci, 277(1694), 2683–2692. 10.1098/rspb.2010.0362

van Heerwaarden, B., & Kellermann, V. (2020). Does plasticity trade off with basal heat tolerance? Trends Ecol Evol, 35(10), 874–885.

van Heerwaarden, B., Kellermann, V., & Sgrò, C. M. (2016). Limited scope for plasticity to increase upper thermal limits. Funct Ecol, 30(12), 1947–1956.

Van Heerwaarden, B., Sgrò, C., & Kellermann, V. M. (2024). Threshold shifts and developmental temperature impact trade-offs between tolerance and plasticity. Proc R Soc B, 291(2016), 20232700. https://pmc.ncbi.nlm.nih.gov/articles/PMC10846935/

Vorhees, A. S., Gray, E. M., & Bradley, T. J. (2013). Thermal resistance and performance correlate with climate in populations of a widespread mosquito. Physiol Biochem Zool, 86(1), 73–81.

Watkins, A. (2021). Testing for phenotypic plasticity. Philos Theor Pract Biol, 13(20220112). 10.3998/ptpbio.16039257.0013.003

West-Eberhard, M. J. (1989). Phenotypic plasticity and the origins of diversity. Annu Rev Ecol System, 249-278.

WHO. (2025). World Health Organization, mosquito-borne diseases. Retrieved July, 15^th^ from https://www.who.int/news-room/fact-sheets/detail/vector-borne-diseases

Wickham, H. (2016). Data analysis : ggplot2 (R package version 3.5.2) [Computer software]. In ggplot2: elegant graphics for data analysis (pp. 189–201). Springer. https://ggplot2.tidyverse.org/

Wickham, H., Averick, M., Bryan, J., Chang, W., McGowan, L. D. A., François, R., Grolemund, G., Hayes, A., Henry, L., & Hester, J. (2019). Welcome to the Tidyverse (R package version 2.0.0). J open source softw, 4(43), 1686.

Wickham, H., François, R., Henry, L., Müller, K., & Vaughan, D. (2023). dplyr: A Grammar of Data Manipulation. (R package version 1.1.4) [Computer software]. In.

Williams, C. R., & Rau, G. (2011). Growth and development performance of the ubiquitous urban mosquito *Aedes notoscriptus* (Diptera: Culicidae) in Australia varies with water type and temperature. Aust J Entomol, 50(2), 195–199.

WMO. (2025). World Meteorological Organization, state of the global climate 2024.. https://library.wmo.int/records/item/69455-state-of-the-global-climate-2024

