## Supplementary information for "Heat hardening enhances mosquito heat tolerance in a species-specific and trait-specific manner"

**Supplementary Data**

Table S1. Pairwise comparisons of estimated LT_50_ values (°C) for post exposure knockdown in *Ae. aegypti*, *Ae. notoscriptus*, and *Cx. quinquefasciatus* following a 1 hr heat shock. Treatment groups: Hardened ♀, Hardened ♂, Basal ♀ and Basal ♂. LT_50_ estimates represent the temperature at which 50% of individuals were knocked down after heat exposure.

|  |  | LT_50_ ratio | Std. Error of the ratio | *t*-value (df=18)  ♀ vs ♂ comparison | *p*-value |
| --- | --- | --- | --- | --- | --- |
| *Ae. aegypti* | |  |  |  |  |
|  | Basal♀/Basal♂ | 1.009 | 0.003 | 3.615 | <0.001 |
|  | Basal♀/Hardened♀ | 0.995 | 0.009 | -2.340 | 0.020 |
|  | Basal♀/ Hardened♂ | 1.006 | 0.002 | 2.950 | 0.003 |
|  | Basal♂/ Hardened♀ | 0.986 | 0.003 | -5.432 | <0.001 |
|  | Basal♂/ Hardened♂ | 0.997 | 0.003 | -1.290 | 0.198 |
|  | Hardened♀/Hardened♂ | 1.011 | 0.002 | 5.172 | <0.001 |
| *Ae. notoscriptus* | |  |  |  |  |
|  | Basal♀/Basal♂ | 1.006 | 0.003 | 2.015 | 0.045 |
|  | Basal♀/Hardened♀ | 0.984 | 0.003 | -5.405 | <0.001 |
|  | Basal♀/ Hardened♂ | 0.992 | 0.003 | -2.366 | 0.019 |
|  | Basal♂/ Hardened♀ | 0.978 | 0.003 | -7.592 | <0.001 |
|  | Basal♂/ Hardened♂ | 0.986 | 0.003 | -4.311 | <0.001 |
|  | Hardened♀/Hardened♂ | 1.009 | 0.003 | 2.607 | 0.010 |
| *Cx. quinquefasciatus* | |  |  |  |  |
|  | Basal♀/Basal♂ | 1.010 | 0.002 | 6.039 | <0.001 |
|  | Basal♀/Hardened♀ | 0.990 | 0.002 | -6.271 | <0.001 |
|  | Basal♀/ Hardened♂ | 1.004 | 0.002 | 2.063 | 0.040 |
|  | Basal♂/ Hardened♀ | 0.980 | 0.001 | -14.400 | <0.001 |
|  | Basal♂/ Hardened♂ | 0.994 | 0.002 | -3.687 | <0.001 |
|  | Hardened♀/Hardened♂ | 1.014 | 0.002 | 8.419 | <0.001 |

Table S2. Pairwise comparisons of estimated LT_50_ values (°C) for 24-hour post-exposure mortality in *Ae. aegypti*, *Ae. notoscriptus*, and *Cx. quinquefasciatus* following a 1 hr heat shock. Treatment groups: Hardened ♀, Hardened ♂, Basal ♀ and Basal ♂. LT_50_ estimates represent the temperature at which 50% mortality occurred 24 hours after heat exposure.

|  |  | LT_50_ ratio | Std. Error of the ratio | *t*-value (df=18) ♀ vs ♂ comparison | *p*-value |
| --- | --- | --- | --- | --- | --- |
| *Ae. aegypti* | |  |  |  |  |
|  | Basal♀/Basal♂ | 1.055 | 0.012 | 4.516 | <0.001 |
|  | Basal♀/Hardened♀ | 0.994 | 0.013 | -0.479 | 0.632 |
|  | Basal♀/ Hardened♂ | 1.054 | 0.011 | 4.768 | <0.001 |
|  | Basal♂/ Hardened♀ | 0.942 | 0.008 | -6.884 | <0.001 |
|  | Basal♂/ Hardened♂ | 0.999 | 0.005 | -0.233 | 0.816 |
|  | Hardened♀/Hardened♂ | 1.060 | 0.008 | 7.270 | <0.001 |
| *Ae. notoscriptus* | |  |  |  |  |
|  | Basal♀/Basal♂ | 1.025 | 0.004 | 6.382 | <0.001 |
|  | Basal♀/Hardened♀ | 0.977 | 0.005 | -4.983 | <0.001 |
|  | Basal♀/ Hardened♂ | 1.002 | 0.003 | 0.768 | 0.443 |
|  | Basal♂/ Hardened♀ | 0.953 | 0.005 | -8.941 | <0.001 |
|  | Basal♂/ Hardened♂ | 0.977 | 0.004 | -5.685 | <0.001 |
|  | Hardened♀/Hardened♂ | 1.026 | 0.005 | 5.149 | <0.001 |
| *Cx. quinquefasciatus* | |  |  |  |  |
|  | Basal♀/Basal♂ | 1.009 | 0.003 | 2.651 | 0.009 |
|  | Basal♀/Hardened♀ | 0.999 | 0.005 | -0.204 | 0.839 |
|  | Basal♀/ Hardened♂ | 1.010 | 0.003 | 2.900 | 0.004 |
|  | Basal♂/ Hardened♀ | 0.990 | 0.004 | -2.393 | 0.018 |
|  | Basal♂/ Hardened♂ | 1.001 | 0.002 | 0.286 | 0.775 |
|  | Hardened♀/Hardened♂ | 1.010 | 0.004 | 2.561 | 0.011 |

Table S3. Type III ANOVA table showing (a) main effects and (b) interaction effects of mosquito species, gender, age and hardening treatment on upper thermal limit (CTmax / ^⸰^C) following a 1 hr heat shock. Factors examined include species (*Ae. aegypti*, *Ae. notoscriptus*, and *Cx. quinquefasciatus*), sex (♀/♂), age group, and prior heat hardening treatment. CTmax represents the critical thermal maximum at which individuals cease movement.

|  | Mean Sq | df | *F-*value | *p-*value |
| --- | --- | --- | --- | --- |
| *Ae. aegypti* |  |  |  |  |
| Main effect model |  |  |  |  |
| Intercept | 25709.9 | 1 | 57222.785 | <0.001 |
| Gender | 4.0 | 1 | 8.912 | 0.003 |
| Age | 12.1 | 1 | 26.994 | <0.001 |
| Hardened | 0.1 | 1 | 0.180 | 0.672 |
| Residuals | 0.4 | 236 |  |  |
| Full interaction model |  |  |  |  |
| Intercept | 7195.8 | 1 | 15815.350 | <0.001 |
| Gender | 0.3 | 1 | 0.552 | 0.458 |
| Age | 2.5 | 1 | 5.556 | 0.019 |
| Hardened | 0.0 | 1 | 0.103 | 0.749 |
| Gender: Age | 0.1 | 1 | 0.117 | 0.732 |
| Gender: Hardened | 0.2 | 1 | 0.351 | 0.554 |
| Age: Hardened | 0.1 | 1 | 0.268 | 0.605 |
| Gender: Age: Hardened | 0.3 | 1 | 0.589 | 0.444 |
| Residuals | 0.5 | 232 |  |  |
| *Ae. notoscriptus* |  |  |  |  |
| Main effect model |  |  |  |  |
| Intercept | 23135.2 | 1 | 152206.091 | <0.001 |
| Gender | 0.2 | 1 | 1.268 | 0.261 |
| Age | 6.2 | 1 | 40.544 | <0.001 |
| Hardened | 0.0 | 1 | 0.215 | 0.643 |
| Residuals | 0.2 | 236 |  |  |
| Full interaction model |  |  |  |  |
| Intercept | 6519.7 | 1 | 42305.047 | <0.001 |
| Gender | 0.0 | 1 | 0.0004 | 0.984 |
| Age | 2.1 | 1 | 13.657 | <0.001 |
| Hardened | 0.0 | 1 | 0.081 | 0.776 |
| Gender: Age | 0.0 | 1 | 0.083 | 0.774 |
| Gender: Hardened | 0.0 | 1 | 0.015 | 0.902 |
| Age: Hardened | 0.0 | 1 | 0.196 | 0.659 |
| Gender: Age: Hardened | 0.0 | 1 | 0.001 | 0.971 |
| Residuals | 0.2 | 232 |  |  |
| *Cx. quinquefasciatus* |  |  |  |  |
| Main effect model |  |  |  |  |
| Intercept | 14257.4 | 1 | 85829.359 | <0.001 |
| Gender | 0.9 | 1 | 5.219 | 0.023 |
| Age | 5.5 | 1 | 33.380 | <0.001 |
| Hardened | 0.2 | 1 | 1.069 | 0.302 |
| Residuals | 0.2 | 231 |  |  |
| Full interaction model |  |  |  |  |
| Intercept | 3801.4 | 1 | 22741.297 | <0.001 |
| Gender | 0.0 | 1 | 0.040 | 0.842 |
| Age | 1.5 | 1 | 8.871 | 0.003 |
| Hardened | 0.1 | 1 | 0.537 | 0.464 |
| Gender: Age | 0.1 | 1 | 0.400 | 0.528 |
| Gender: Hardened | 0.0 | 1 | 0.037 | 0.847 |
| Age: Hardened | 0.1 | 1 | 0.641 | 0.424 |
| Gender: Age: Hardened | 0.0 | 1 | 0.003 | 0.955 |
| Residuals | 0.2 | 227 |  |  |

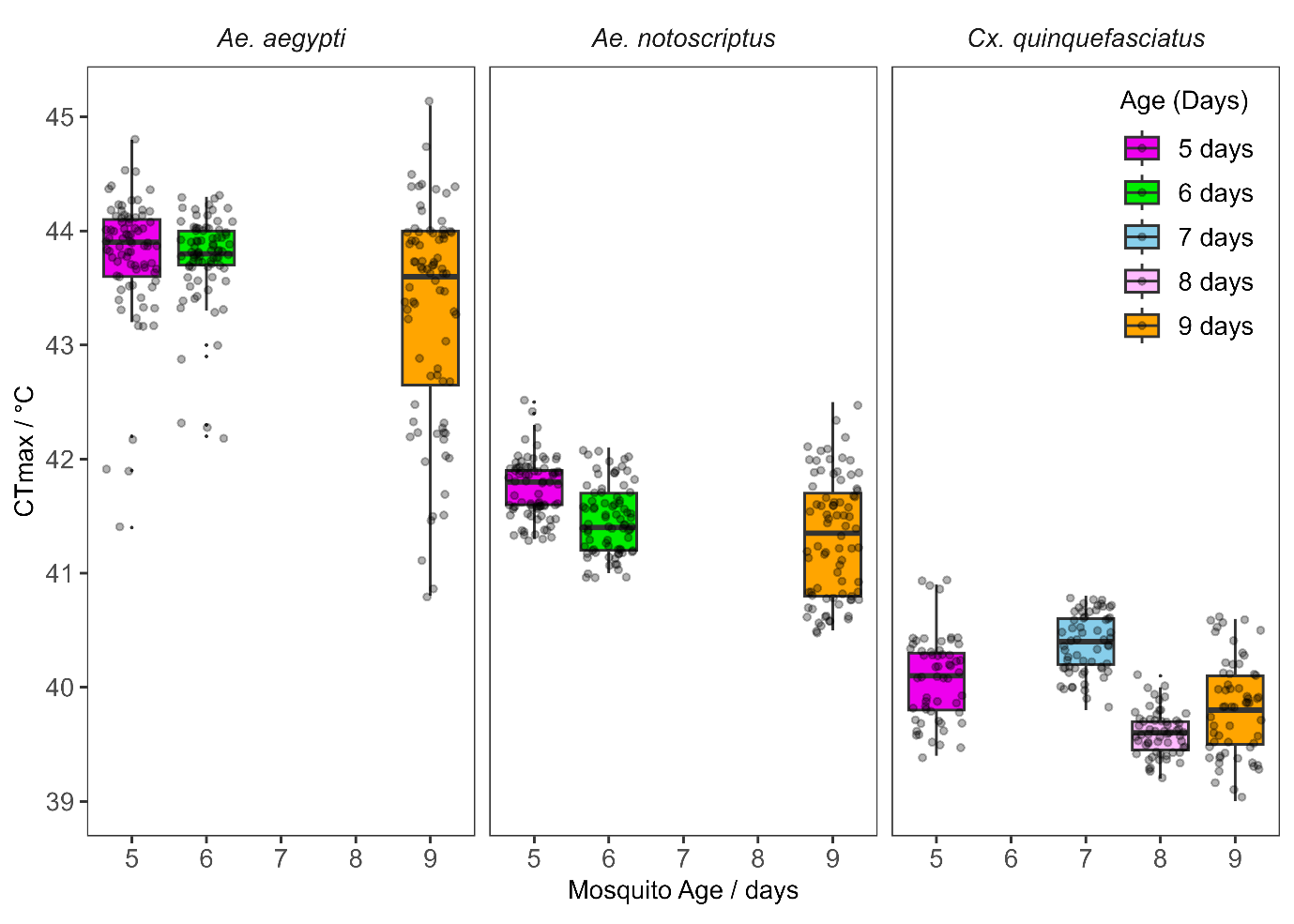

Figure S1. Box plot for effect of age/block on the CTmax of *Ae. aegypti*, *Ae. notoscriptus*, and *Cx. quinquefasciatus* mosquitoes across 3-4 age groups. Each boxplot includes both sexes and hardening treatments for a given age/block. CTmax represents the critical thermal maximum at which individuals cease movement. Horizontal lines show medians, boxes represent 25^th^ and 75^th^ percentiles and whiskers represent the minimum and the maximum values excluding outliers.

Table S4. Type III ANOVA table showing (a) main effects and (b) interaction effects of sex, age and hardening treatment on heat knockdown time (minutes) in *Ae. aegypti*, *Ae. notoscriptus*, and *Cx. quinquefasciatus* following a 1 hr heat shock. Heat knockdown time refers to the duration until movement ceased under thermal stress.

|  | Mean Sq | df | *F-*value | *p-*value |
| --- | --- | --- | --- | --- |
| *Ae. aegypti* |  |  |  |  |
| Main effect model |  |  |  |  |
| Intercept | 6841.7 | 1 | 375.969 | <0.001 |
| Gender | 36.3 | 1 | 1.992 | 0.159 |
| Age | 82.2 | 1 | 4.519 | 0.035 |
| Hardened | 55.8 | 1 | 3.067 | 0.081 |
| Residuals | 18.2 | 236 |  |  |
| Full interaction model |  |  |  |  |
| Intercept | 2507.8 | 1 | 137.676 | <0.001 |
| Gender | 15.5 | 1 | 0.849 | 0.358 |
| Age | 124.0 | 1 | 6.805 | 0.010 |
| Hardened | 79.0 | 1 | 4.334 | 0.038 |
| Gender: Age | 28.4 | 1 | 1.558 | 0.213 |
| Gender: Hardened | 27.3 | 1 | 1.501 | 0.222 |
| Age: Hardened | 64.1 | 1 | 3.519 | 0.062 |
| Gender: Age: Hardened | 32.0 | 1 | 1.756 | 0.186 |
| Residuals | 18.2 | 232 |  |  |
| *Ae. notoscriptus* |  |  |  |  |
| Main effect model |  |  |  |  |
| Intercept | 5924.6 | 1 | 494.641 | <0.001 |
| Gender | 9.3 | 1 | 0.778 | 0.379 |
| Age | 168.6 | 1 | 14.074 | <0.001 |
| Hardened | 14.6 | 1 | 1.223 | 0.270 |
| Residuals | 12.0 | 236 |  |  |
| Full interaction model |  |  |  |  |
| Intercept | 1434.89 | 1 | 119.586 | <0.001 |
| Gender | 4.55 | 1 | 0.379 | 0.539 |
| Age | 8.98 | 1 | 0.749 | 0.388 |
| Hardened | 2.29 | 1 | 0.191 | 0.663 |
| Gender: Age | 12.11 | 1 | 1.009 | 0.316 |
| Gender: Hardened | 0.66 | 1 | 0.055 | 0.815 |
| Age: Hardened | 2.56 | 1 | 0.213 | 0.645 |
| Gender: Age: Hardened | 0.04 | 1 | 0.003 | 0.956 |
| Residuals | 12.00 | 232 |  |  |
| *Cx. quinquefasciatus* |  |  |  |  |
| Main effect model |  |  |  |  |
| Intercept | 586.3 | 1 | 36.811 | <0.001 |
| Gender | 213.7 | 1 | 13.415 | <0.001 |
| Age | 889.5 | 1 | 55.846 | <0.001 |
| Hardened | 22.3 | 1 | 1.398 | 0.238 |
| Residuals | 15.9 | 234 |  |  |
| Full interaction model |  |  |  |  |
| Intercept | 279.3 | 1 | 17.745 | <0.001 |
| Gender | 61.6 | 1 | 3.914 | 0.049 |
| Age | 102.0 | 1 | 6.484 | 0.012 |
| Hardened | 0.3 | 1 | 0.021 | 0.884 |
| Gender: Age | 45.7 | 1 | 2.905 | 0.090 |
| Gender: Hardened | 0.0 | 1 | 0.0001 | 0.993 |
| Age: Hardened | 0.9 | 1 | 0.059 | 0.808 |
| Gender: Age: Hardened | 1.6 | 1 | 0.102 | 0.750 |
| Residuals | 15.7 | 230 |  |  |

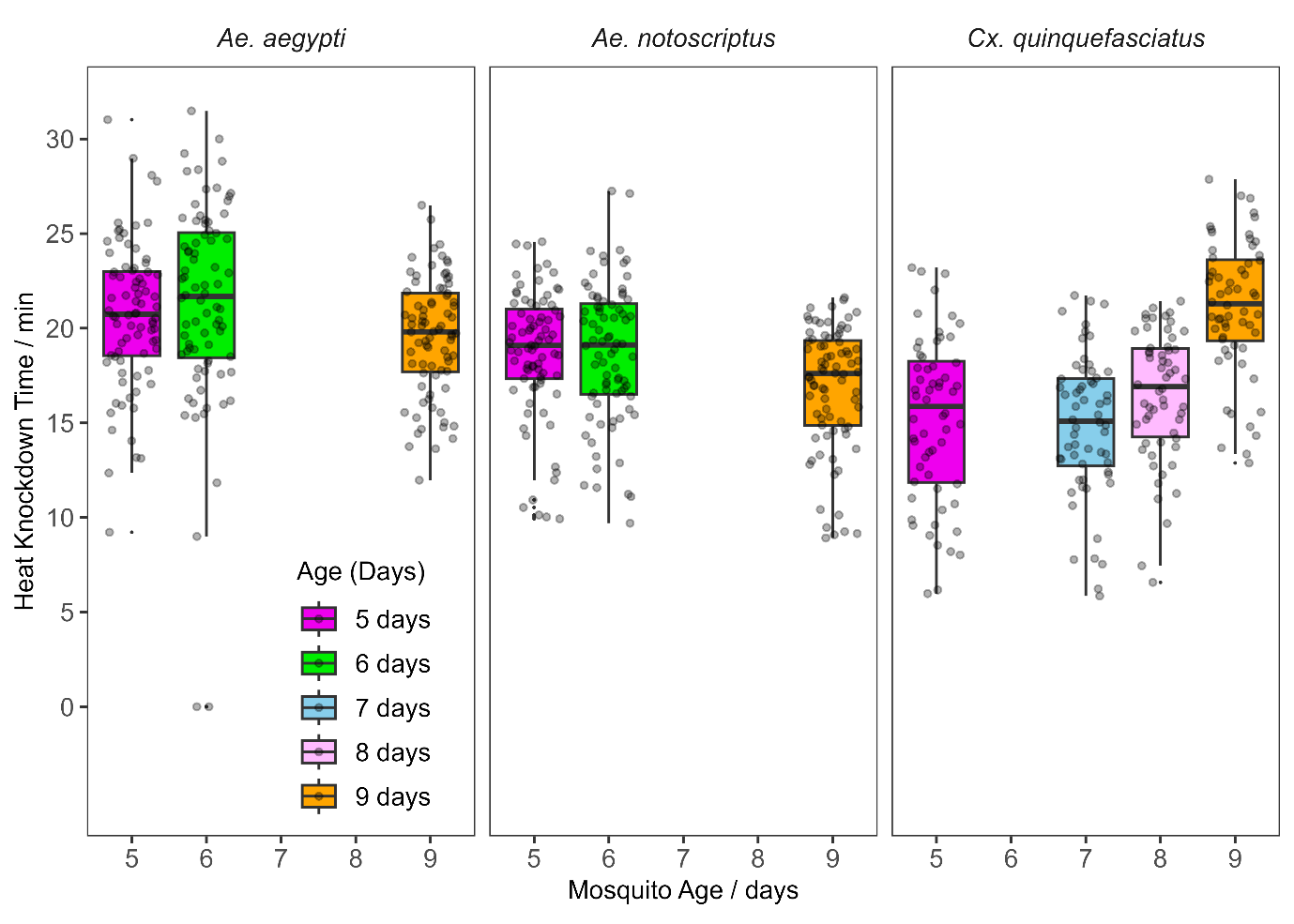

Figure S2. Box plot for effect of age/block on the heat knockdown time of *Ae. aegypti* (A), *Ae. notoscriptus* (B), and *Cx. quinquefasciatus* (C) mosquitoes across 3-4 age groups. Knockdown time was tested at 43.1 °C for *Ae. aegypti*, 41.2 °C for *Ae. notoscriptus*, and 39.8°C for *Cx. quinquefasciatus.* Each boxplot includes both sexes and hardening treatments for a given age/block. Heat knockdown time represents the duration until individuals ceased movement under acute thermal stress. Horizontal lines show medians, boxes represent 25^th^ and 75^th^ percentiles and whiskers represent the minimum and the maximum values excluding outliers.

Table S5. Pairwise comparison of estimated LT_50_ values (°C) for post exposure knockdown and 24-hour post exposure mortality in *Ae. aegypti* offspring, following a 1 hr heat shock in the parental generation. Treatment groups: Treated parent’s offspring ♀, Treated parent’s offspring ♂, Untreated parent’s offspring ♀ and Untreated parent’s offspring ♂. LT_50_ values represent the temperature at which 50% of individuals were knocked down or died.

|  |  | LT_50_ ratio | Std. Error of the ratio | *t*-value (df=18) ♀ vs ♂ comparison | *p*-value |
| --- | --- | --- | --- | --- | --- |
| Knockdown | |  |  |  |  |
|  | Untreated ♀ parents’ offspring ♀/ Untreated ♀ parents’ offspring ♂ | 1.003 | 0.002 | 1.839 | 0.067 |
|  | Untreated ♀ parents’ offspring ♀/ Treated ♀ parents’ offspring ♀ | 1.003 | 0.002 | 1.526 | 0.128 |
|  | Untreated ♀ parents’ offspring ♀/ Treated ♀ parents’ offspring ♂ | 1.011 | 0.002 | 6.180 | <0.001 |
|  | Untreated ♀ parents’ offspring ♂/ Treated ♀ parents’ offspring ♀ | 0.999 | 0.002 | -0.324 | 0.746 |
|  | Untreated ♀ parents’ offspring ♂/ Treated ♀ parents’ offspring ♂ | 1.008 | 0.002 | 4.430 | <0.001 |
|  | Treated ♀ parents’ offspring ♀/ Treated ♀parents’ offspring ♂ | 1.009 | 0.002 | 4.759 | <0.001 |
| Mortality | |  |  |  |  |
|  | Untreated ♀ parent’s offspring ♀/ Untreated ♀ parents’ offspring ♂ | 1.015 | 0.004 | 3.984 | <0.001 |
|  | Untreated ♀ parents’ offspring ♀/ Treated ♀ parents’ offspring ♀ | 0.989 | 0.009 | -1.233 | 0.219 |
|  | Untreated ♀ parents’ offspring ♀/ Treated ♀ parents’ offspring ♂ | 1.019 | 0.003 | 6.638 | <0.001 |
|  | Untreated ♀ parents’ offspring ♂/ Treated ♀ parents’ offspring ♀ | 0.974 | 0.010 | -2.748 | 0.006 |
|  | Untreated ♀ parents’ offspring ♂/ Treated ♀ parents’ offspring ♂ | 1.004 | 0.004 | 0.973 | 0.331 |
|  | Treated ♀ parents’ offspring ♀/ Treated ♀ parents’ offspring ♂ | 1.031 | 0.010 | 3.171 | 0.002 |

Table S6. Type III ANOVA table showing (a) main effects and (b) interaction effects of *Ae. aegypti* offspring sex and parental heat exposure (1 hr heat shock at 41^0^C) on upper thermal limit (CTmax / ^0^C) and heat knockdown time (minutes). CTmax represents the critical thermal maximum at which individuals ceased movement, while heat knockdown time refers to the duration until movement ceased under thermal stress.

|  | Mean Sq | df | *F*-value | *p-*value |
| --- | --- | --- | --- | --- |
| Upper thermal limit (CTmax) |  |  |  |  |
| Main effect model |  |  |  |  |
| Intercept | 153966.0 | 1 | 1196809.870 | <0.001 |
| Gender | 0.0 | 1 | 1.411 | 0.236 |
| Parental exposure | 0.0 | 1 | 0.802 | 0.372 |
| Residuals | 0.1 | 236 |  |  |
| Full interaction model |  |  |  |  |
| Intercept | 115475.0 | 1 | 900577.071 | <0.001 |
| Gender | 0.0 | 1 | 3.185 | 0.076 |
| Parental exposure | 0.0 | 1 | 0.094 | 0.760 |
| Gender: Parental exposure | 0.0 | 1 | 1.780 | 0.183 |
| Residuals | 0.1 | 235 |  |  |
| Heat knockdown Time |  |  |  |  |
| Main effect model |  |  |  |  |
| Intercept | 11935.0 | 1 | 1067.525 | <0.001 |
| Gender | 0.1 | 1 | 0.005 | 0.944 |
| Parental exposure | 0.8 | 1 | 0.074 | 0.786 |
| Residuals | 11.2 | 237 |  |  |
| Full interaction model |  |  |  |  |
| Intercept | 8981.7 | 1 | 800.008 | <0.001 |
| Gender | 0.2 | 1 | 0.014 | 0.907 |
| Parental exposure | 0.8 | 1 | 0.067 | 0.795 |
| Gender: Parental exposure | 0.1 | 1 | 0.009 | 0.924 |
| Residuals | 11.2 | 236 |  |  |
